# Computational identification of candidate human long noncoding RNAs associated with PBMC responses to *Candida auris* and its cell wall components

**DOI:** 10.64898/2026.09.21.753102

**Authors:** Md. Takim Sarker, Bledar Bisha

**Affiliations:** Department of Animal Science, University of Wyoming, Laramie, United States

**Author notes:** Corresponding author: Bledar Bisha.

**Keywords:** *Candida auris*, long noncoding RNA, peripheral blood mononuclear cells, host-pathogen interaction, innate immunity, RNA sequencing

## Abstract

*Candida auris* is an emerging multidrug-resistant fungal pathogen. However, little is known about the role of long noncoding RNAs (lncRNAs) in the human immune response to this fungus. This study aimed to identify and prioritize novel lncRNAs associated with the response of human peripheral blood mononuclear cells (PBMCs) to C. auris and its cell wall components. We reanalyzed RNA-sequencing data from peripheral blood mononuclear cells obtained from three healthy donors. The cells were exposed to live *C. auris*, purified mannan, purified β-glucan, or RPMI control for 4 h or 24 h. We identified 607 putative novel polyadenylated lncRNA loci that were absent from GENCODE Release 48. Of the 449 novel loci tested for differential expression, 102 responded to at least one stimulus. Novel-lncRNA responses were limited at 4 h but increased markedly at 24 h. At 24 h, 78 novel lncRNAs responded to live *C. auris*, 34 to mannan, and 11 to β-glucan. Many of these responses are specific to a particular stimulus. Among the 102 responsive lncRNAs, 94 were significantly coexpressed with protein-coding genes. The coding genes of the most highly connected lncRNAs were associated with innate immunity, defense responses, lysosomal functions, and signaling pathways. Temporal analysis revealed 19 novel lncRNAs whose responses changed between 4 and 24 h. Fourteen novel lncRNAs were also differentially expressed and were selected for further analysis. Combining temporal response, coexpression, and genomic proximity identified 4 candidate cis-associated pairs: MSTRG.32011-TLR2, MSTRG.5763-DUSP5, MSTRG.25627-HCK, and MSTRG.18159-ITGB3. These findings reveal a predominantly late and stimulus-specific novel-lncRNA response to *C. auris* and provide prioritized candidates for experimental validation.

## Introduction

*Candida auris* is an emerging fungal pathogen that causes invasive infections and healthcare-associated outbreaks. Whole-genome analysis revealed that genetically different and often antifungal-resistant lineages emerged at nearly the same time on several continents ^1^. Its capacity to remain viable on healthcare surfaces further facilitates persistence and transmission within clinical environments ^2^. These characteristics distinguish *C. auris* from many other Candida species and emphasize the need to understand the host responses that influence infection and clearance.

Interactions between *C. auris* and the innate immune system are complex. Compared with *C. albicans* neutrophils, human neutrophils exhibit limited recruitment, extracellular trap formation, and killing in response to *C. auris* ^3^. However, studies using human peripheral blood mononuclear cells (PBMCs) have demonstrated substantial cytokine production and transcriptional changes following exposure to *C. auris* ^4^. These responses depend partly on fungal cell wall composition and the duration of exposure. In particular, β-glucan contributes strongly to the early PBMC response, whereas mannan has a greater influence at later time points ^4^. Differences in cell wall structure have also been associated with variations in the magnitude of innate immune activation ^5^. Together, these findings indicate that the host response to *C. auris* is dynamic and depends on the immune cell population, fungal stimulus, and time after exposure.

Most transcriptomic studies of antifungal immunity have focused on protein-coding genes. Long noncoding RNAs (lncRNAs), which are conventionally defined as transcripts of at least 200 nucleotides with limited protein-coding potential, represent an additional and incompletely characterized layer of gene regulation. Compared with protein-coding genes, human lncRNAs frequently show greater tissue and context specificity, allowing previously unannotated transcripts to emerge under particular biological conditions ^6^. Experimental studies have established that individual lncRNAs can influence innate immune transcription. For example, lincRNA-Cox2 regulates both the activation and repression of immune response genes following pattern recognition receptor stimulation ^7^, whereas lincRNA-EPS restrains inflammatory gene expression in macrophages ^8^. These findings provide a biological basis for investigating previously unannotated lncRNAs during fungal stimulation, such as in *C. auris*.

The contribution of lncRNAs to antifungal host responses remains poorly understood. Host lncRNA expression has been examined during experimental *C. albicans* infection in mice ^9^, whereas studies involving *C. auris* have investigated mainly lncRNAs encoded by the fungus itself ^10, 11^. The available human PBMC transcriptomic studies of *C. auris* have focused primarily on protein-coding responses and have not systematically identified previously unannotated human lncRNA loci ^4^. To our knowledge, novel human lncRNA responses to live *C. auris* and its purified cell wall components have therefore not been comprehensively characterized.

In this study, we reanalyzed donor-matched 3′ RNA-sequencing data from human PBMCs exposed to live *C. auris*, purified *C. auris* mannan, purified *C. auris* β-glucan, or medium control (RPMI) medium for 4 h and 24 h. We sought to identify previously unannotated polyadenylated lncRNA candidates and determine whether their expression varied according to stimulus and time. Differential expression, coexpression, functional enrichment, temporal interaction, genomic proximity, and network analyses were subsequently integrated to prioritize lncRNAs for experimental investigation. This approach was designed to extend the established protein-coding transcriptional response and provide a focused catalog of candidate human lncRNAs associated with the immune response to *C. auris*.

## Methods

### Study design and RNA-sequencing dataset

This study comprised a secondary analysis of publicly available RNA-sequencing data generated by Bruno *et al*. to investigate human immune responses to *Candida auris* ^4^. Raw sequencing data were obtained from NCBI BioProject PRJNA647871. The present analysis was restricted to 24 samples representing three healthy human donors, four stimulation conditions, and two time points **(Supplementary Table S1)**. The stimulation conditions were as follows: RPMI medium control, live *C. auris* strain KCTC17810, purified *C. auris* mannan, and purified *C. auris* β-glucan. Each donor contributed one sample to every stimulus-time combination, producing three biological replicates for each of the eight experimental groups.

As described in the original study ^4^, peripheral blood mononuclear cells (PBMCs) were isolated from venous blood obtained from healthy donors and cultured at 5 × 10⁶ cells mL^-1^. The cells were incubated in RPMI containing 10% pooled human serum and stimulated with live *C. auris* at 1 × 10⁶ cells mL^-1^, mannan at 10 µg mL^-1^, β-glucan at 10 µg mL^-1^, or medium alone for 4 h or 24 h. QuantSeq 3′ mRNA-Seq FWD libraries were prepared using 100 ng of total RNA from donors A and 250 ng from donors B and C and sequenced as 75-bp single-end reads on an Illumina NextSeq 500. Because the RNA input and library preparation were donor-associated, the donor was included explicitly in all differential expression models.

### Overview of the bioinformatic workflow

The complete computational workflow is summarized in **Figure 1**. The RNA-sequencing reads were quality controlled, trimmed, aligned and assembled before the candidate transcripts were subjected to structural, noncoding potential, expression support and manual review filters. The resulting novel-lncRNA annotation was then used for differential expression, overlapping, coexpression, enrichment, temporal interaction, cis-proximity and network analyses.

**Figure 1.**
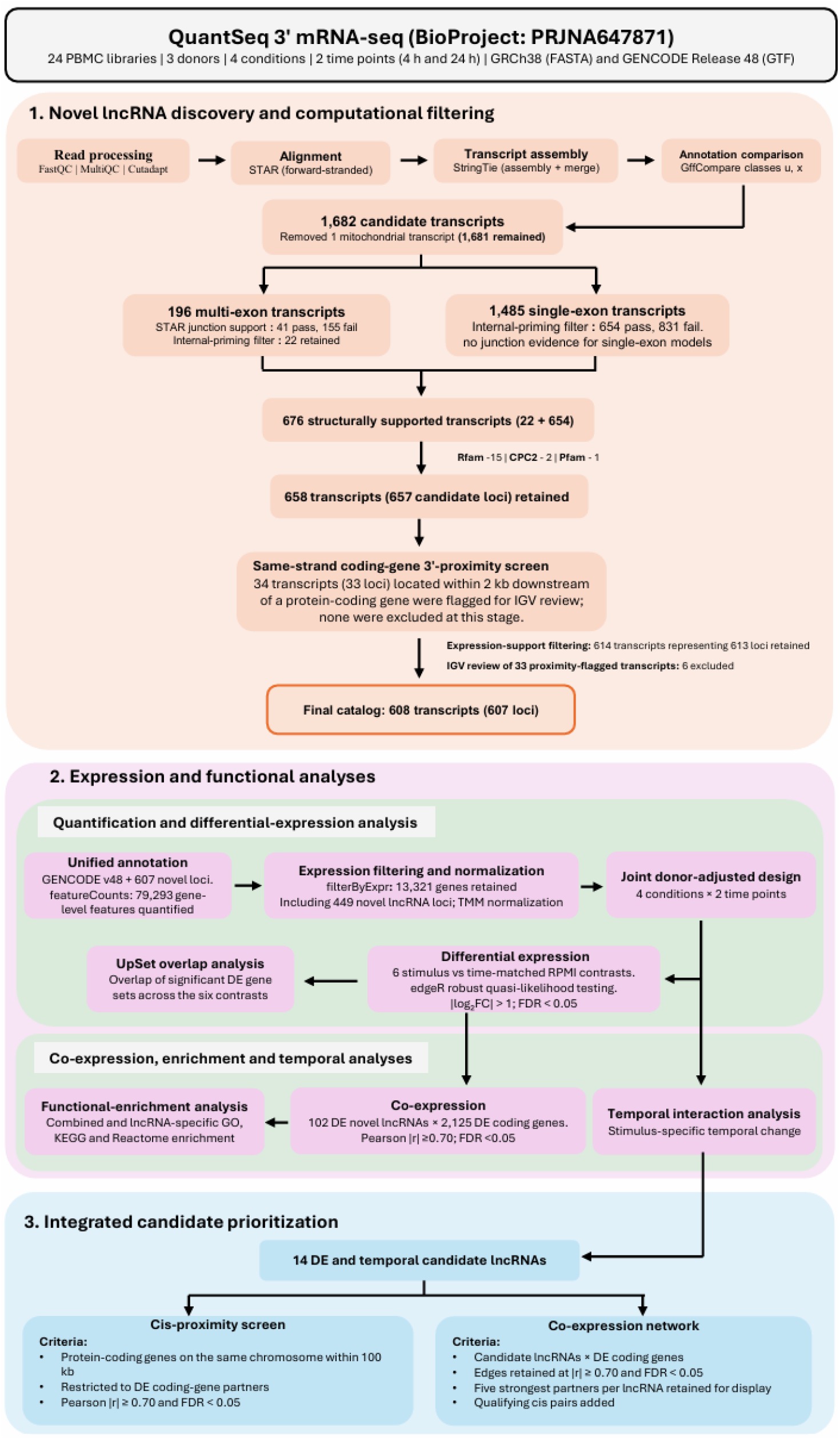
Bioinformatic workflow for novel lncRNA discovery and analysis. The workflow summarizes RNA-sequencing processing, transcript assembly, candidate lncRNA filtering and the downstream analyses used to characterize and prioritize novel lncRNAs.

### Reference genome and gene annotation

The human reference genome fasta file [Genome sequence, primary assembly (GRCh38)] and annotation GTF file (Comprehensive gene annotation, CHR) were obtained from GENCODE Release 48, which is based on GRCh38.p14 ^12^. Reference FASTA and GTF files were restricted to the 25 reference chromosomes shared between the files (chromosomes 1-22, X, Y, and mitochondrial chromosome M) to maintain consistent sequence identifiers. A STAR v2.7.11b ^13^ genome index was generated with a splice junction database overhang of 74 nucleotides, corresponding to the 75-bp read length.

### Read preprocessing, quality control, and alignment

Raw and trimmed reads were assessed with FastQC v0.12.1 ^14^. The reports were aggregated across the 24 samples via MultiQC v1.35 ^15^. Reads were processed with Cutadapt v5.2 ^16^ via a three-stage pipeline designed for QuantSeq libraries. First, terminal poly(A) and poly(G) sequences containing at least 20 consecutive bases were removed, with a minimum retained read length of 20 nucleotides and a maximum of two adapter-removal rounds. Second, NextSeq quality trimming was applied at a cutoff of 10, followed by removal of the QuantSeq 3′ adapter sequence A{18}AGATCGGAAGAGCACACGTCTGAACTCCAGTCAC with a minimum overlap of three nucleotides and a maximum error rate of 0.1. Third, reads containing an extended match of at least 20 nucleotides to AGATCGGAAGAGCACACGTCTGAACTCCAGTCAC at the 5′ end were discarded. Reads shorter than 20 nucleotides after any stage were excluded. The trimmed reads were aligned to the human reference via STAR v2.7.11b ^13^. The alignment was performed in two-pass basic mode, with coordinate-sorted BAM output, intron-motif strand annotation, and gene-count output enabled. BAM files were indexed with SAMtools 1.23.1 ^17^.

### Novel lncRNA discovery and computational filtering

#### Transcriptome assembly and identification of novel transcript candidates

Transcript-guided assembly was performed independently for each sample via StringTie v3.0.3 ^18^. Assemblies were generated using the forward-stranded library specification, GENCODE Release 48 as the reference annotation, a minimum assembled transcript length of 200 nucleotides, and eight processing threads. The expression-only assembly mode was not used, allowing StringTie to report transcript structures absent from the reference annotation. The 24 sample-level assemblies were combined with StringTie merge, using the same reference annotation and minimum transcript length. The merged assembly was compared with GENCODE via GffCompare v0.12.10 ^19^. Transcripts assigned class code u, denoting intergenic transcripts, or x, denoting exonic overlap with a reference transcript on the opposite strand, were retained as primary novel candidates. Class-code “i” transcripts were excluded because they could not be reliably distinguished from retained introns or unprocessed precursor RNA. Mitochondrial candidates were also excluded. Candidate sequences of at least 200 nucleotides were extracted with GffRead v0.12.9 ^19^.

Because 3′ QuantSeq is not designed for complete transcript reconstruction, the retained features were interpreted as candidate novel polyadenylated lncRNA loci rather than definitive full-length transcript isoforms.

#### Splice-junction validation and internal-priming filtering

Each intron in a multiexon candidate was matched by genomic position and strand using the STAR SJ.out.tab files from all 24 samples. An intron was accepted when it had a recognized STAR splice motif (codes 1-6), was supported by uniquely mapped reads in at least two samples and by at least three reads across all samples and had a maximum junction overhang of at least 12 nt. Multiexon transcripts were retained only when every intron passed these criteria; this validation was not applicable to single-exon transcripts. To identify potential internal oligo(dT)-priming artifacts, the 20 genomic nucleotides immediately downstream of each candidate’s 3′ end were extracted in transcript orientation. Candidates were classified as A-rich when this sequence contained either ≥6 consecutive adenines or ≥12 adenines overall ^20^. Only non-A-rich candidates were advanced for subsequent analyses.

#### Structural-RNA and coding-potential filters

Candidate sequences were screened against Rfam release 15.1 (retrieved 30 July 2026) via cmscan from INFERNAL 1.1.5 (Sep 2023) ^21,22^. The searches used Rfam curated gathering thresholds, clan information, top-strand-only analysis, and the --rfam, --nohmmonly, and --cut_ga options. Candidates with matches to recognized structural RNA families, including ribosomal RNA, transfer RNA, or signal-recognition-particle RNA, were excluded.

Protein-coding potential was assessed with CPC2 v1.0.1 ^23^. Candidates classified as coding were removed. Candidate open reading frames were then identified with orfipy v0.0.4 ^24^. Forward-strand ORFs of at least 30 nucleotides beginning with ATG were reported via genetic code 1; partial 3′ ORFs were permitted because of the 3′-enriched library design. The longest predicted ORF from each transcript was translated and searched against Pfam-A release 38.2 via hmmscan from HMMER v3.4 and family-specific gathering thresholds ^25,26^. Candidates containing a significant Pfam domain were excluded.

#### Assessment of possible protein-coding gene readthroughs

To identify candidate signals that might originate from annotated coding genes, 2-kb windows were constructed immediately downstream of the 3′ end of each GENCODE protein-coding gene via strand-aware genomic operations implemented with BEDTools v2.31.1 ^27^. Candidate loci overlapping these windows on the same strand were marked as having elevated readthrough or terminal-exon extension risk. This screen was used as a review flag rather than an automatic exclusion criterion.

#### Expression support and manual genomic review

A provisional annotation file containing GENCODE Release 48 and all lncRNA candidates surviving the sequence-based filters was constructed. Gene-level read counts were generated from all 24 BAM files with featureCounts v2.1.1 using exons as counting features, gene_id as the grouping attribute, and forward-strand counting (-s 1) ^28^. Multimapping reads were not enabled for counting. Next, for a candidate locus (lncRNA) to receive expression support, it was required to have counts per million (CPM) ≥ 0.5 in at least two different donors belonging to the same stimulus-time group. This criterion requires reproducible expression within a biologically matched condition and prevents support from being assigned solely on the basis of repeated samples from one donor. Expression-supported candidates carrying a same-strand 3′-proximity flag were inspected via the Integrative Genomics Viewer (IGV v2.19.8) ^29^.

In the IGV, candidate and GENCODE annotations, strand-specific alignments, splice junctions, and read coverage were examined across all donors. Candidates were retained when they displayed a reproducible, spatially distinct transcriptional signal separated from the neighboring protein-coding gene by a coverage gap. Candidates showing continuous readthrough coverage from the coding gene or evidence that they represented an unannotated terminal-exon extension were excluded. Ambiguous cases and the reasons for each exclusion were recorded in **Supplementary Table S2**.

The retained novel lncRNA transcript models were appended to the complete GENCODE Release 48 annotation. The resulting unified annotation was used with featureCounts to quantify all 24 aligned libraries at the gene level.

### Quantification and differential expression analysis

#### Gene-level quantification, classification, and expression filtering

Genes were classified as protein coding, known lncRNA, novel lncRNA, or other transcript types. The operational known-lncRNA category included GENCODE records annotated as lncRNA, lincRNA, antisense, processed_transcript, sense_intronic, or sense_overlapping. Novel candidates were recognized from their assigned novel_lncRNA annotation or original MSTRG identifier. Statistical analyses were performed in R v4.5.1 via edgeR v4.6.3 ^30^. Lowly expressed genes were filtered with filterByExpr, and the eight stimulus-time combinations were used as the experimental groups. Library size normalization factors were estimated via the trimmed mean of M values (TMM) method **(Supplementary Figure S1A)** ^31^. For exploratory visualization, log2-CPM values were calculated with a prior count of 2, and principal component analysis was performed without additional feature scaling **(Supplementary Figure S1B)**.

#### Differential expression analysis

All 24 samples were analyzed jointly via the design ∼0 + group + donor, where the group represented the eight combinations of stimulation conditions and time points **(Table 1)**, and the donor was included as a fixed blocking factor. Negative-binomial dispersions were estimated via estimateDisp(…, robust=TRUE), followed by quasi likelihood model fitting with glmQLFit(…, robust=TRUE) ^32^. Six prespecified contrasts compared live *C. auris*, mannan, and β-glucan with time-matched RPMI controls at 4 h and 24 h via glmQLFTest. P values were adjusted across all tested genes within each contrast via the Benjamini-Hochberg procedure. Differential expression was defined as an FDR < 0.05 and a |log₂-fold change | > 1.

**Table 1.** Experimental groups and temporal interaction contrasts. Each stimulation condition was evaluated at 4 h and 24 h, producing eight stimulus-time groups. Temporal interactions were used to determine whether the stimulus effect was related to the time-matched RPMI control, which differed between 4 h and 24 h. A positive interaction coefficient indicates a greater stimulus effect at 24 h, whereas a negative coefficient indicates a reduced or more negative effect at 24 h.

| Stimulation<br>condition | 4 h group | 24 h group | Temporal interaction contrast |
| --- | --- | --- | --- |
| RPMI control | RPMI<sub>4h</sub> | RPMI<sub>24h</sub> | Reference condition |
| Live *C. auris* | Live<sub>4h</sub> | Live<sub>24h</sub> | (Live<sub>24h</sub> - RPMI<sub>24h</sub>) - (Live<sub>4h</sub> - RPMI<sub>4h</sub>) |
| *C. auris* mannan | Mannan<sub>4h</sub> | Mannan<sub>24h</sub> | (Mannan<sub>24h</sub> - RPMI<sub>24h</sub>) - (Mannan<sub>4h</sub> - RPMI<sub>4h</sub>) |
| *C. auris* β-glucan | β-glucan<sub>4h</sub> | β-glucan<sub>24h</sub> | (β-glucan<sub>24h</sub> - RPMI<sub>24h</sub>) - (β-glucan<sub>4h</sub> - RPMI<sub>4h</sub>) |

#### Overlap of differential expression results across stimuli and time points

Gene sets meeting the differential expression criteria (FDR < 0.05 and | log₂-fold change | > 1) were compared across the six stimulus-versus-RPMI contrasts at 4 h and 24 h. Shared and contrast-specific gene sets were visualized via the R package UpSetR ^33^. Comparisons were performed for all differentially expressed genes and separately for novel lncRNAs, both irrespective of expression direction and separately for upregulated and downregulated genes.

### Coexpression, enrichment and temporal analyses

#### lncRNA-protein coding gene coexpression analysis

Coexpression analysis was restricted to novel lncRNAs and protein-coding genes that were differentially expressed in at least one primary stimulus contrast. The log2-CPM values with a prior count of 2 were adjusted for the donor via limma::removeBatchEffect ^34^ while preserving the eight-level stimulus-time design. Pearson correlations were calculated across the 24 residualized sample profiles for every novel lncRNA-protein coding gene pair.

Correlation P values were calculated from the Pearson t statistic using 20 residual degrees of freedom, and Benjamini-Hochberg correction was applied jointly across all tested pairs. Significant associations were defined by |r| ≥ 0.70 and FDR < 0.05. Because the observations represented repeated experimental measurements from three donors, this analysis was used for exploratory prioritization; coexpression was not interpreted as evidence of direct regulatory relationships.

The novel lncRNAs were ranked by the number of significant protein-coding partners. The 15 highest degree lncRNAs were used for the global coexpression heatmap and functional enrichment analyses. For heatmap display, protein-coding genes were ranked first by the number of significant associations with these 15 lncRNAs and then by the mean absolute correlation; the 25 highest-ranked genes are displayed.

#### Functional enrichment analysis

Functional enrichment was performed on the union of significant protein-coding partners of the 15 highest-degree novel lncRNAs. The background universe comprised all the differentially expressed protein-coding genes included in the coexpression analysis rather than the complete genome.

Gene symbols were converted to Entrez identifiers via org.Hs.eg.db v3.21.0. ^35^ Gene Ontology biological process enrichment was performed via clusterProfiler v4.16.0 ^36^, with gene set sizes restricted to 10-500 genes, Benjamini-Hochberg-adjusted P < 0.05, and q < 0.20. Reactome enrichment was performed with ReactomePA v1.52.0 via analogous gene set limits, adjusted P < 0.05, and q < 0.05 ^37^. For KEGG pathway enrichment, the human organism database was used, with gene set sizes ranging from 10-500 and adjusted P < 0.05 and q < 0.05 ^38^.

Secondary enrichment analyses were performed separately for the coding partners of individual high-degree lncRNAs. Because multiplicity correction was performed independently within each lncRNA analysis, these lncRNA-specific enrichments were treated as exploratory.

### Temporal interaction analysis

Stimulus-specific temporal changes were assessed with difference-in-differences contrasts of the form (stimulus24h - RPMI24h) - (stimulus4h - RPMI4h) **(Table 1)**.

Separate interaction contrasts were evaluated for live *C. auris*, mannan, and β-glucan via the same donor-adjusted quasi likelihood model. Temporal interactions were considered significant at FDR < 0.05, and the absolute interaction log2-fold change was > 1. Novel lncRNAs prioritized as stimulus-responsive and time-dependent were required to be differentially expressed in at least one same-time stimulus contrast and to exhibit a significant interaction in response to at least one stimulus.

### Cis-proximity and integrated network analyses

#### Genomic proximity and identification of candidate cis associations

Each temporally prioritized novel lncRNA was compared with GENCODE protein-coding genes on the same chromosome. Genomic distance was defined as the interval separating the two gene bodies, with overlapping intervals assigned a distance of zero. Pairs separated by no more than 100 kb were classified as proximal. A proximal pair was designated a high-confidence exploratory cis association only when the protein-coding gene was differentially expressed and the lncRNA-coding-gene correlation satisfied |r| ≥ 0.70 and FDR < 0.05. Proximity and coexpression were not interpreted as demonstrating direct cis regulation.

#### Network construction and visualization

The complete set of significant coding gene associations involving the temporally prioritized novel lncRNAs was exported as node and edge tables. Edges were annotated with the Pearson correlation coefficient, adjusted P value, correlation direction, genomic distance, and cis-proximity classification.

For a reduced visualization network, the five most significant coding gene partners of each lncRNA were retained after being ranked by FDR and then by absolute correlation. All high-confidence proximal pairs were added regardless of rank. The resulting network was visualized via Cytoscape v3.10.4 ^39^. The network degree represents the number of statistical coexpression associations and was not interpreted as the degree of molecular regulation.

### Use of generative AI and AI-assisted technologies

During the preparation of this manuscript and its accompanying computational materials, the authors used OpenAI ChatGPT and Codex to assist with paraphrasing and restructuring author-provided text, language editing, formatting, code debugging, and documentation of selected analysis and visualization code. The authors reviewed and revised all the AI-assisted text and independently checked the code, numerical results, references, figures, interpretations, and conclusions against the underlying data and original sources. The AI tools did not generate experimental data or make final scientific decisions. The authors take full responsibility for the accuracy and integrity of the submitted work.

## Results

### Sequencing quality and read retention after trimming

Across the 24 raw libraries, MultiQC reported 129.80 million reads **(Supplementary Data 1**). The reads averaged approximately 74 nucleotides (nt) in length and presented consistently high base-call quality (Q34-Q36) with negligible ambiguous base content. Adapter contamination was detected in most libraries prior to trimming.

After Cutadapt processing, 129.05 million reads were retained (**Supplementary Data 2**). Only 0.75 million reads (0.58%) were removed. The mean read length decreased slightly from 73.93 to 71.72 nt, indicating that trimming primarily removed short adapter or poly(A)-contaminated terminal sequences. Following trimming, all 24 libraries passed per-base quality and adapter-content assessments, with no evidence of excessive loss of reads.

### Reference-genome alignment performance

The mean overall mapping rate was 99.22%, ranging from 98.57% to 99.45% (**Supplementary Data 3**). The mean percentage of uniquely mapped reads was 80.42%, with a range of 75.93-83.99%. The mean mismatch rate was 0.54% and varied only from 0.51% to 0.56%, while the insertion and deletion rates were each approximately 0.01%. Across the libraries, 99.68% of the detected splice events matched junctions present in the reference annotation. Inspection of the strand-specific STAR gene-count columns revealed substantially greater read assignment in the forward orientation than in the reverse orientation, supporting the expected forward-strand library configuration.

### Identification and filtering of candidate novel lncRNAs

Reference-guided StringTie assemblies from the 24 libraries were merged and compared with the GENCODE Release 48 annotation GTF files via GffCompare. The merged assembly contained 400,205 transcript models across 79,978 loci. Of these, 384,464 were exact matches to reference transcripts (class code =), whereas 388 were classified as intergenic (u) and 1,294 showed exonic overlap with an annotated transcript on the opposite strand (x). The 1,682 u and x transcript models were selected as the primary discovery set (**Supplementary Table S3).** After one mitochondrial transcript was removed, 1,681 nuclear candidates remained, including 196 multiexon and 1,485 single-exon transcripts. All 1,681 candidates satisfied the minimum spliced-length requirement of 200 nucleotides **(Table 2)**.

**Table 2.**
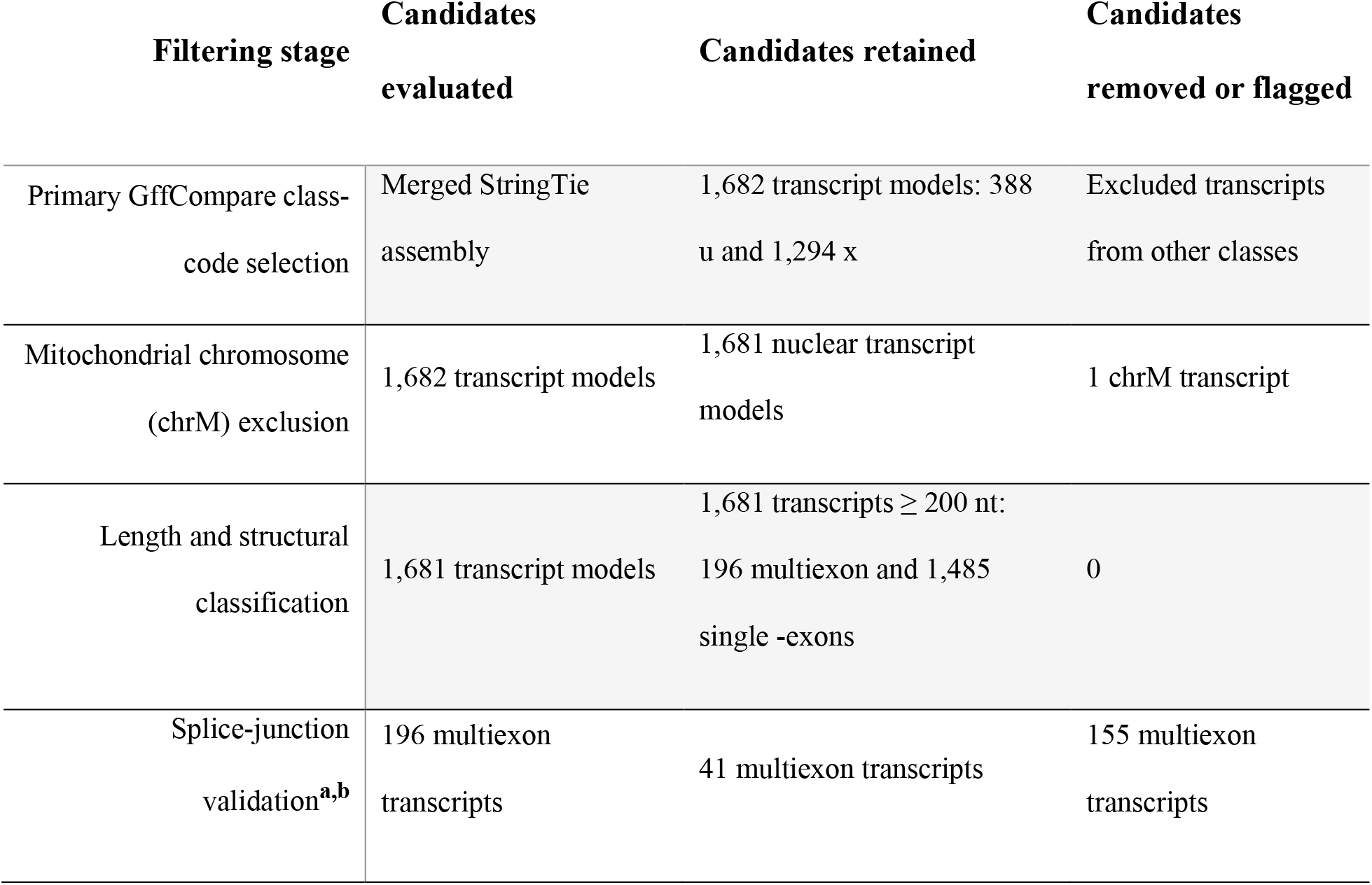
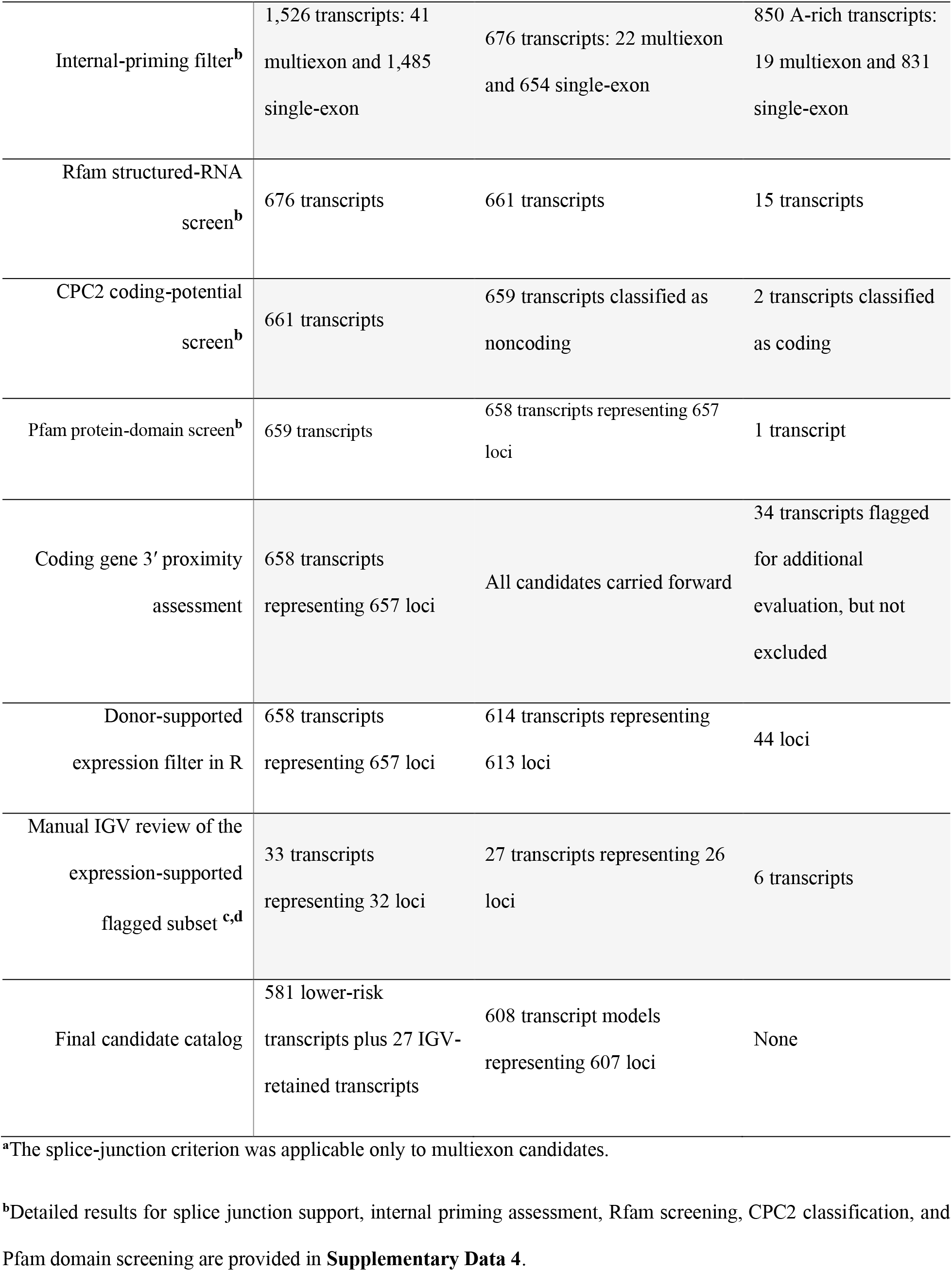
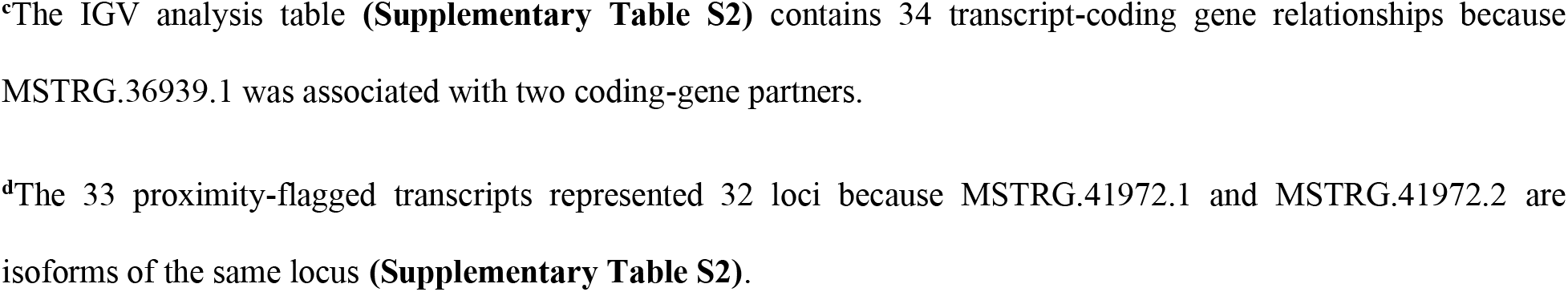
Sequential filtering of candidate novel lncRNA transcript models.

| Filtering stage | Candidates evaluated | Candidates retained | Candidates removed or flagged |
| --- | --- | --- | --- |
| Primary GffCompare class-code selection | Merged StringTie assembly | 1,682 transcript models: 388 u and 1,294 x | Excluded transcripts from other classes |
| Mitochondrial chromosome (chrM) exclusion | 1,682 transcript models | 1,681 nuclear transcript models | 1 chrM transcript |
| Length and structural classification | 1,681 transcript models | 1,681 transcripts ≥ 200 nt:<br>196 multiexon and 1,485 single -exons | 0 |
| Splice-junction validation<sup>a,b</sup> | 196 multiexon transcripts | 41 multiexon transcripts | 155 multiexon transcripts |
| Internal-priming filter<sup>b</sup> | 1,526 transcripts: 41 multiexon and 1,485 single-exon | 676 transcripts: 22 multiexon and 654 single-exon | 850 A-rich transcripts: 19 multiexon and 831 single-exon |
| Rfam structured-RNA screen<sup>b</sup> | 676 transcripts | 661 transcripts | 15 transcripts |
| CPC2 coding-potential screen<sup>b</sup> | 661 transcripts | 659 transcripts classified as noncoding | 2 transcripts classified as coding |
| Pfam protein-domain screen<sup>b</sup> | 659 transcripts | 658 transcripts representing 657 loci | 1 transcript |
| Coding gene 3' proximity assessment | 658 transcripts representing 657 loci | All candidates carried forward | 34 transcripts flagged for additional evaluation, but not excluded |
| Donor-supported expression filter in R | 658 transcripts representing 657 loci | 614 transcripts representing 613 loci | 44 loci |
| Manual IGV review of the expression-supported flagged subset <sup>c,d</sup> | 33 transcripts representing 32 loci | 27 transcripts representing 26 loci | 6 transcripts |
| Final candidate catalog | 581 lower-risk transcripts plus 27 IGV-retained transcripts | 608 transcript models representing 607 loci | None |

<sup>a</sup>The splice-junction criterion was applicable only to multiexon candidates.
<sup>b</sup>Detailed results for splice junction support, internal priming assessment, Rfam screening, CPC2 classification, and Pfam domain screening are provided in **Supplementary Data 4**.
<sup>c</sup>The IGV analysis table (**Supplementary Table S2**) contains 34 transcript-coding gene relationships because MSTRG.36939.1 was associated with two coding-gene partners.
<sup>d</sup>The 33 proximity-flagged transcripts represented 32 loci because MSTRG.41972.1 and MSTRG.41972.2 are isoforms of the same locus (**Supplementary Table S2**).

Among the 196 multiexon candidates, 41 had all introns supported by the required STAR splice junction evidence, whereas 155 were excluded. Internal-priming analysis subsequently revealed 22 of these 41 multiexon candidates and 654 of the 1,485 single-exon candidates. The resulting structural candidate set therefore contained 676 transcript models.

Rfam analysis revealed 49 significant matches distributed among 15 candidates. These matches represented structured RNA families, including ribosomal RNA, transfer RNA, and signal-recognition-particle RNA, and the corresponding 15 candidates were excluded. Among the remaining 661 transcripts, CPC2 classified 659 as noncoding and two as coding. Pfam analysis identified a protein-domain match in one additional candidate. This produced 658 putative lncRNA transcript models representing 657 genomic loci.

Screening for possible same-strand extensions of annotated protein-coding genes identified 34 transcript models located within the predefined 2-kb region downstream of an annotated coding-gene 3′ end. These models were flagged for further evaluation in the IGV but were not removed at this stage. Next, expression screening revealed that 613 of the 657 loci (93.3%) **(Figure 2A, B)** corresponded to 614 transcript models, whereas 44 loci failed to reach the requirement of CPM ≥ 0.5 in at least two donors within the same stimulus-time group **(Supplementary Table S4).**

**Figure 2.**
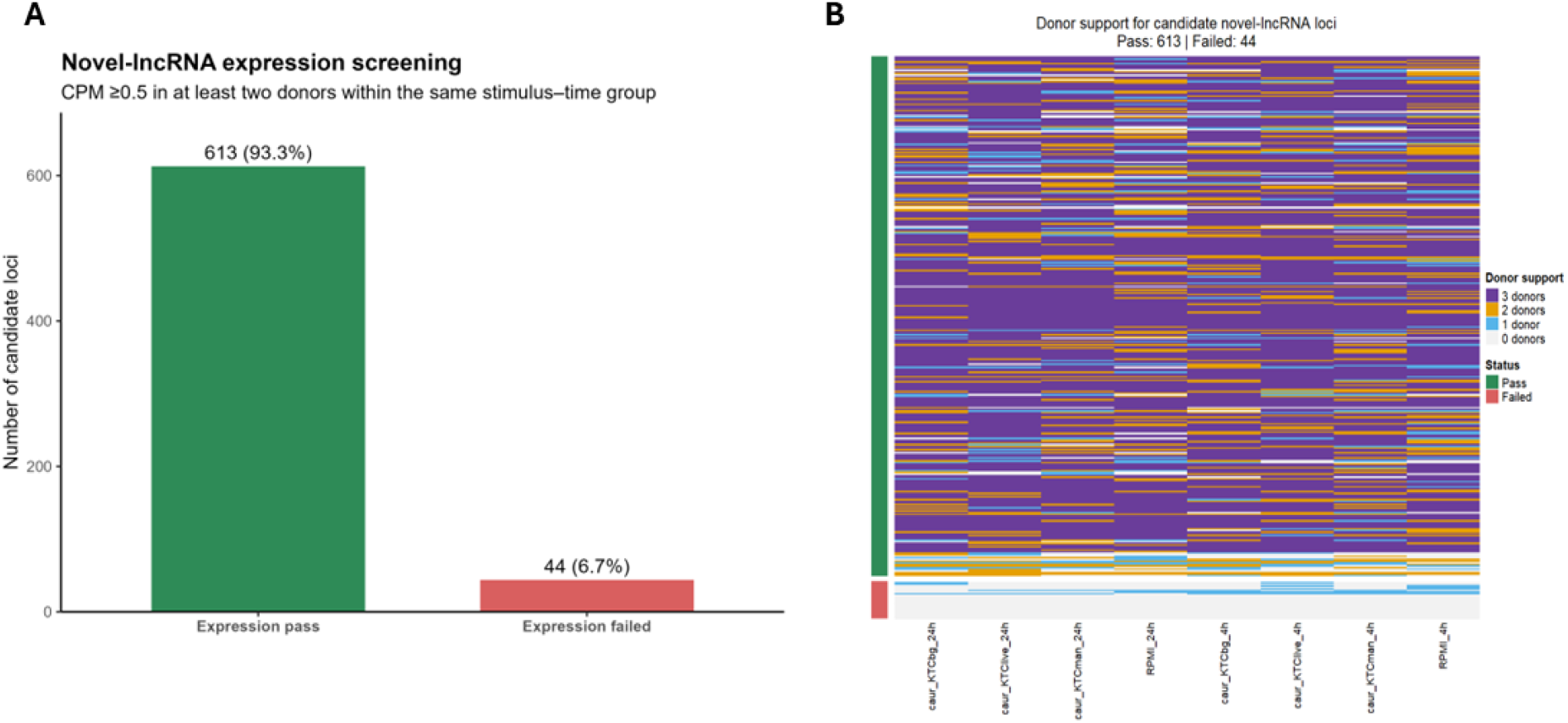
Donor-supported expression filtering of candidate novel lncRNA loci. **(A)** Numbers and proportions of candidate loci passing or failing the expression-support criterion. A locus passed when it reached CPM ≥ 0.5 in at least two of the three donors within at least one stimulus-time group. Among the 657 candidate loci evaluated, 613 (93.3%) passed, and 44 (6.7%) failed. **(B)** Heatmap showing the number of supporting donors for each candidate locus across the eight stimulus-time groups formed by RPMI, β-glucan, live *Candida auris*, and mannan at 4 h and 24 h.

Among the expression-supported candidates, 33 of the same-strand 3′-proximity flagged transcript models, representing 32 loci, were subjected to manual IGV assessment **(Supplementary Table S2)**. Six genes were excluded because their coverage was more consistent with the extension of a neighboring protein-coding gene. Twenty-seven flagged transcript models representing 26 loci were retained. Combining these 27 manually accepted models with 581 expression-supported, lower-risk models produced a final catalog of 608 transcript models representing 607 putative novel polyadenylated lncRNA loci. Joint quantification via the complete GENCODE Release 48 annotation and the final novel-lncRNA catalog produced a unified count matrix containing 78,686 GENCODE loci and 607 putative novel lncRNA loci across the 24 libraries.

### Comparatively more genes were differentially expressed at 24 h than at 4 h

After count-based expression filtering, 13,321 genes were retained for differential expression testing. These genes included 11,173 protein-coding genes, 1,434 annotated lncRNAs, 449 of the 607 novel lncRNA loci, and 265 genes belonging to other biotypes.

PCA separated the 4 h and 24 h samples along PC1 (37.3% of variance), whereas PC2 (13.3%) distinguished live-*C. auris*, mannan, and β-glucan-stimulated samples from RPMI controls at 24 h **(Supplementary Figure S1B)**. RPMI samples are also separated by time, highlighting the importance of time-matched controls.

The transcriptional responses were limited at 4 h but expanded substantially by 24 h. At 4 h, β-glucan, live *C. auris*, and mannan stimulation produced 262, 193, and 93 differentially expressed genes, respectively. At 24 h, the corresponding numbers increased to 509, 1,933, and 1,208 genes, respectively. Therefore, live *C. auris* produced the broadest late response, followed by mannan and β-glucan **(Table 3**; **Figure 3A).** The complete differential expression results for all six stimulus versus time-matched RPMI contrasts are provided in **Supplementary Data 5.**

**Figure 3.**
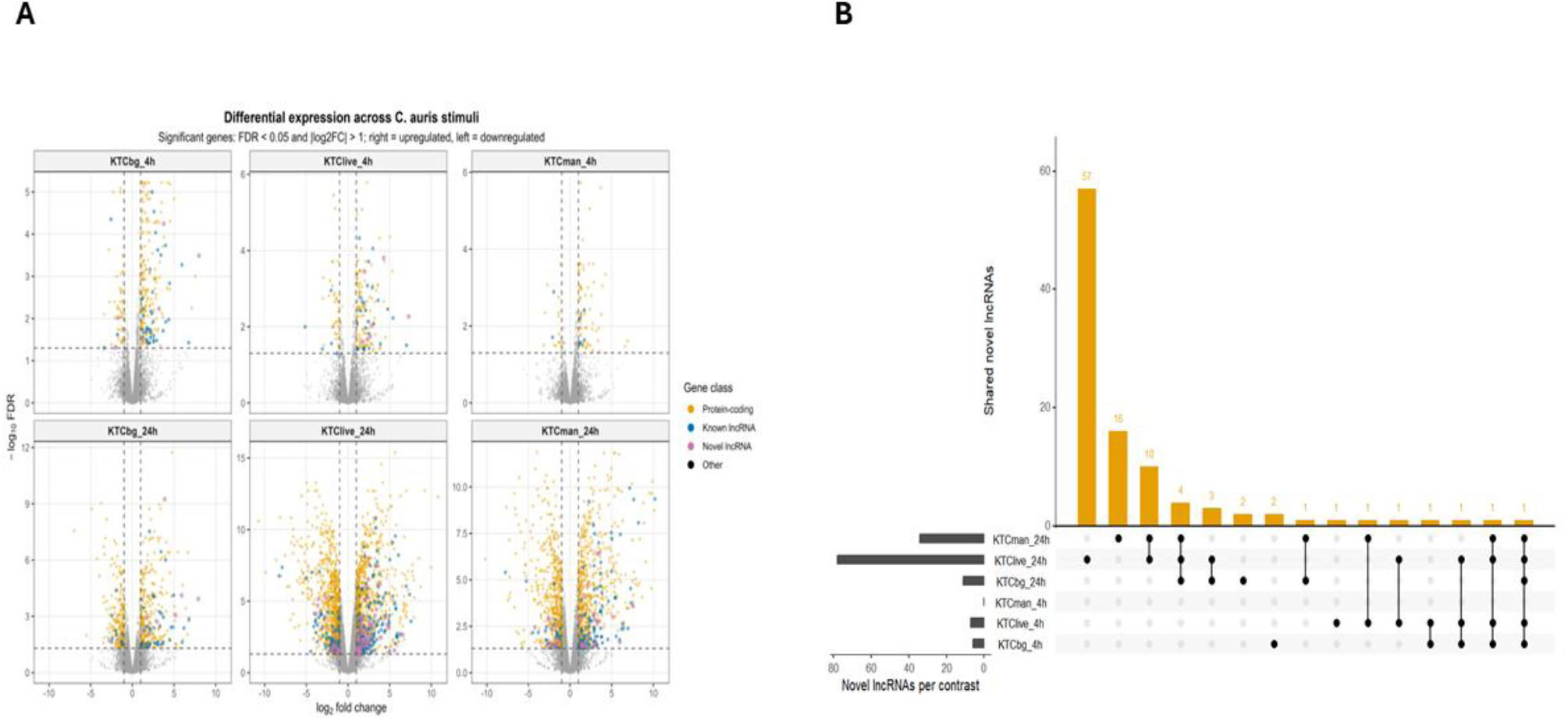
Differential expression patterns and overlap of novel lncRNAs across stimulus-time contrasts. (A) Volcano plots showing differential expression of β-glucan, live *Candida auris*, and mannan-stimulated PBMCs relative to the corresponding time-matched RPMI controls at 4 h and 24 h. The x-axis represents the log2-fold change, and the y-axis represents the - log10 false discovery rate (FDR); genes are distinguished according to the annotation class. Differential expression was defined as an FDR < 0.05 and a |log2FC| > 1. **(B)** UpSet plot showing the overlap among the 102 unique differentially expressed novel lncRNAs across the six contrasts, irrespective of expression direction. The horizontal bars indicate the number of novel lncRNAs in each contrast, the vertical bars indicate intersection sizes, and the connected dots identify the contrasts contributing to each intersection.

**Table 3.** Differentially expressed genes in each stimulus-time comparison.

| Contrast | Genes Tested | Up regulated | Down regulated | Total DE | Protein coding DE | Known lncRNA DE | Novel lncRNA DE |
| --- | --- | --- | --- | --- | --- | --- | --- |
| KTCbg_4h | 13321 | 211 | 51 | 262 | 214 | 40 | 6 |
| KTClive_4h | 13321 | 148 | 45 | 193 | 155 | 29 | 7 |
| KTCman_4h | 13321 | 75 | 18 | 93 | 85 | 8 | 0 |
| KTCbg_24h | 13321 | 311 | 198 | 509 | 446 | 52 | 11 |
| KTClive_24h | 13321 | 998 | 935 | 1933 | 1601 | 240 | 78 |
| KTCman_24h | 13321 | 692 | 516 | 1208 | 1039 | 128 | 34 |
| Sum of contrast specific DE calls<sup>a</sup> | 13321 | 2435 | 1763 | 4198 | 3540 | 497 | 136 |

<sup>a</sup>The same 13,321 genes were tested in each comparison. The values in the “Sum of contrast-specific DE calls” row (except the 2<sup>nd</sup> column) represent summed contrast-specific DE calls rather than unique genes; a gene significant in multiple contrasts is counted once in each corresponding contrast. Therefore, the 136 novel-lncRNA DE calls represent 102 unique novel lncRNAs.

The novel lncRNAs presented a similar time-dependent pattern. At 4 h, 6 novel lncRNAs responded to β-glucan, 7 responded to live *C. auris*, and none responded to mannan. At 24 h, these numbers increased to 11, 78, and 34, respectively. In total, 102 unique novel lncRNAs were differentially expressed in at least one stimulus-time comparison. These results indicate that most detectable novel-lncRNA responses developed during the later phase of stimulation.

Overlap analysis revealed that a substantial proportion of the novel-lncRNA response was stimulus-specific **(Figure 3B)**. Among the 78 novel lncRNAs detected following live-*C. auris* stimulation at 24 h, 57 were unique to that contrast. Sixteen of the 34 novel lncRNAs that responded to mannan at 24 h were unique to mannan, while the largest shared intersection contained ten novel lncRNAs detected following both live-*C. auris* and mannan stimulation at 24 h. Direction-specific overlap patterns for up and downregulated novel lncRNAs are presented in **Supplementary Figure S2**.

### Coexpression analysis identified highly connected novel lncRNAs

Across the six contrasts **(Table 3),** 3,540 protein-coding DE calls represented 2,125 unique genes, whereas 136 novel-lncRNA DE calls represented 102 unique loci. Testing all 216,750 possible lncRNA-coding gene pairs revealed 10,866 significant associations involving 94 novel lncRNAs and 1,774 protein-coding genes. Eight differentially expressed novel lncRNAs had no coding gene partners meeting these criteria.

The 15 most highly connected lncRNAs had between 260 and 440 significant coding gene partners. MSTRG.2194 presented the highest connectivity, with 440 partners, followed by MSTRG.39337 and MSTRG.22879, with 410 and 408 partners, respectively. A heatmap of these 15 lncRNAs and 25 selected coding genes illustrated contrasting positive and negative coexpression patterns **(Figure 4A).** These highly connected lncRNAs were selected for subsequent functional enrichment analysis.

**Figure 4.**
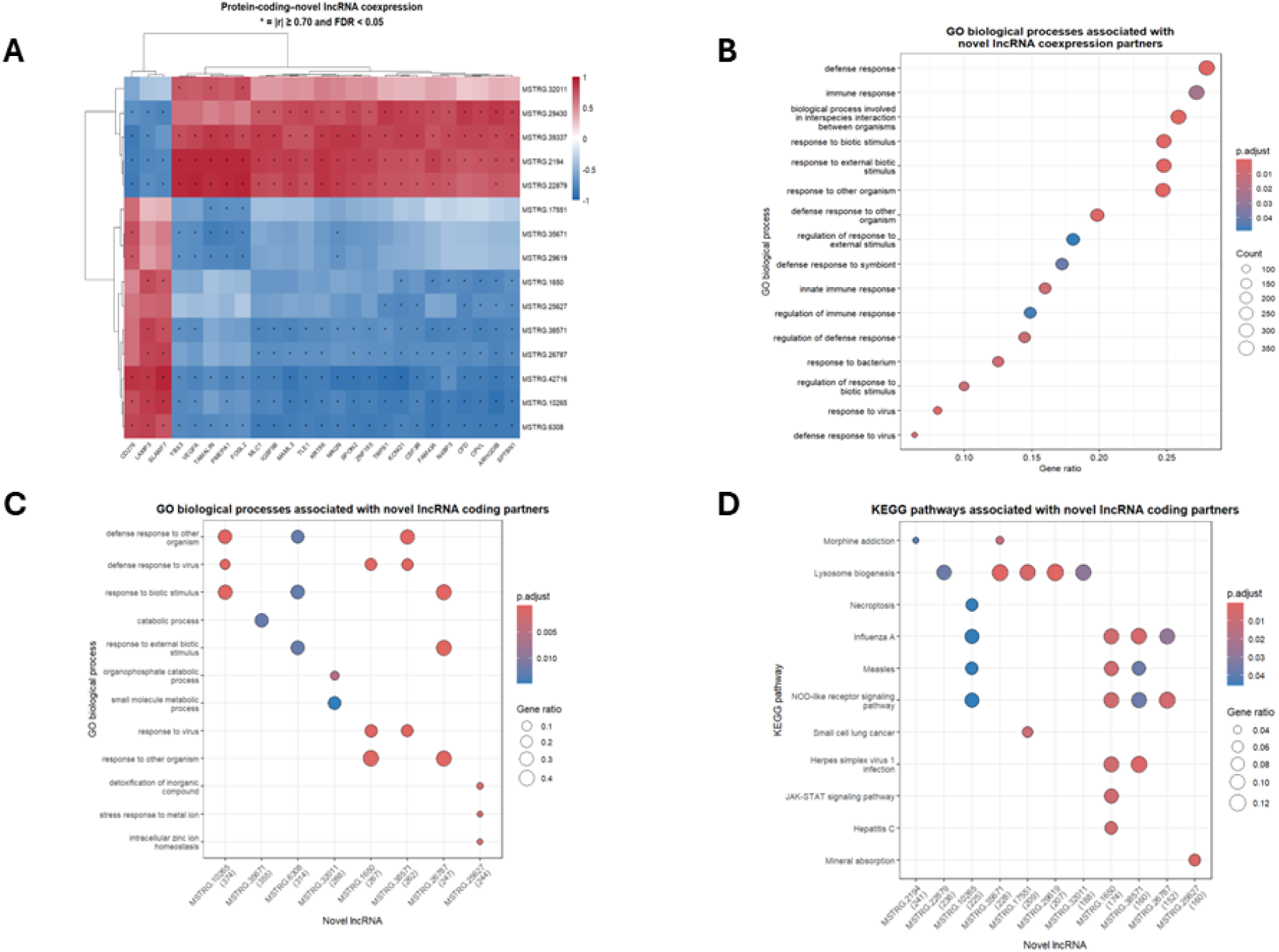
Coexpression and functional enrichment of highly connected novel lncRNAs. **(A)** Correlation heatmap of 15 highly connected novel lncRNAs and 25 selected coding genes. Red and blue indicate positive and negative correlations, respectively; asterisks indicate |r| ≥ 0.70 and FDR < 0.05. **(B)** GO enrichment of their combined coding gene partners. **(C–D)** GO and KEGG enrichment of individual lncRNA partner sets. Dot color indicates BH-adjusted P values; dot size represents gene counts in **B** and gene ratios in **C-D**. The parenthetical counts in **C-D** indicate database-annotated partners used as gene ratio denominators.

### Functional enrichment among coding gene partners of highly connected novel lncRNAs

The 15 highest-connectivity lncRNAs collectively had 1,370 distinct protein-coding partners. Enrichment analysis of this combined partner set revealed 16 significant Gene Ontology (GO) biological process terms, including response to biotic stimulus, defense response, innate immune response, and regulation of immune response **(Figure 4B)**. No KEGG or Reactome pathways met the significance criteria in the combined-set analysis.

Analyses of individual lncRNA partner sets revealed significant GO enrichment for 8 of the 15 lncRNAs and significant KEGG enrichment for 11 **[Figure 4(C, D)]**. The KEGG results revealed 25 lncRNA-pathway associations involving 11 distinct pathways. Lysosome biogenesis was enriched among the partners of five lncRNAs, whereas NOD-like receptor signaling was enriched among the partners of four lncRNAs. Other findings included JAK-STAT signaling among MSTRG.1650 partners and mineral absorption among MSTRG.25627 partners **(Supplementary Table S5)**. These results associate the selected lncRNAs with coding gene sets enriched for immune-related, lysosomal, and metabolic functions without establishing that the lncRNAs regulate those processes.

### Temporal interaction analysis identified time-dependent novel lncRNA responses

Temporal interaction analysis revealed that 178 genes responded differently between 4 h and 24 h following β-glucan stimulation, 945 following live *C. auris* stimulation, and 646 following mannan stimulation. Among these genes, 2, 10, and 10 genes were classified as novel lncRNAs **(Supplementary Table S6)**. These 22 stimulus-specific interactions involved 19 distinct novel lncRNA loci, with 3 loci showing significant temporal interactions under 2 stimulation conditions.

None of the 22 stimulus-lncRNA combinations met the single-time-point differential expression criteria at 4 h. At 24 h, 10 were upregulated, 6 were downregulated, and 6 remained below the differential expression threshold. At the locus level, 14 of the 19 novel lncRNAs were also identified by primary differential expression analysis, comprising 9 late-induced and 5 late-repressed candidates. The remaining 5 loci were identified only by the temporal interaction test.

Three loci showed significant temporal interactions with two stimuli. MSTRG.18159 exhibited positive interactions following both β-glucan and live *C. auris* stimulation, with interaction log2 fold changes of 6.18 and 5.06, respectively. MSTRG.2547 was expressed late following live *C. auris* and mannan stimulation, whereas MSTRG.2229 exhibited negative interactions with β-glucan and live *C. auris* without meeting the single-time-point differential expression criteria.

### Genomic proximity and coexpression support candidate cis associations

The 14 novel lncRNAs meeting both the differential expression criterion and the temporal interaction criterion were evaluated for proximity to annotated protein-coding genes. Initial same-chromosome matching generated 14,930 lncRNA-coding gene pairs. Restricting these genes to a gene-body distance of ≤100 kb resulted in the retention of 60 proximal pairs, involving 13 novel lncRNAs and 56 distinct protein-coding genes **(Supplementary Table S7)**.

Among these 60 pairs, 13 involved coding genes belonging to the previously identified set of 2,125 differentially expressed protein-coding genes. The remaining 47 pairs included genes outside this set. The 13 eligible pairs represented 12 distinct coding genes. A comparison with the existing coexpression results revealed that 4 pairs met both association criteria, |r| ≥ 0.70 and FDR < 0.05, whereas 9 did not.

The 4 supported pairs were MSTRG.32011-TLR2, MSTRG.5763-DUSP5, MSTRG.25627-HCK, and MSTRG.18159-ITGB3 **(Table 4)**. All the correlations were positive, ranging from 0.747 to 0.937.

**Table 4.**
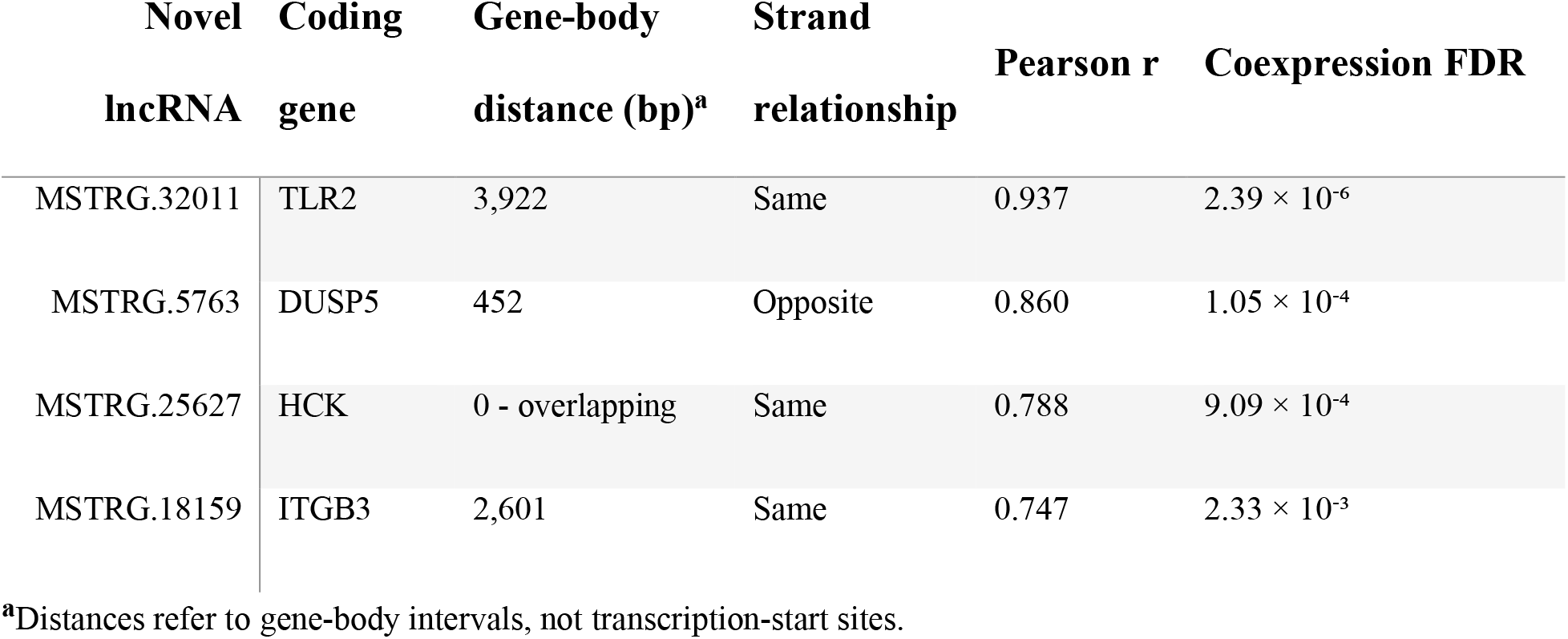
Prioritized proximal lncRNA-protein coding gene associations.

| Novel<br>lncRNA | Coding<br>gene | Gene-body<br>distance (bp)<sup>a</sup> | Strand<br>relationship | Pearson r | Coexpression FDR |
| --- | --- | --- | --- | --- | --- |
| MSTRG.32011 | TLR2 | 3,922 | Same | 0.937 | $2.39 \times 10^{-6}$ |
| MSTRG.5763 | DUSP5 | 452 | Opposite | 0.860 | $1.05 \times 10^{-4}$ |
| MSTRG.25627 | HCK | 0 - overlapping | Same | 0.788 | $9.09 \times 10^{-4}$ |
| MSTRG.18159 | ITGB3 | 2,601 | Same | 0.747 | $2.33 \times 10^{-3}$ |

<sup>a</sup>Distances refer to gene-body intervals, not transcription-start sites.

MSTRG.32011-TLR2 exhibited the strongest correlation and was separated by 3,922 bp, whereas MSTRG.25627 overlapped the HCK gene-body interval. Among these 4 lncRNAs, MSTRG.32011 and MSTRG.25627 also belonged to the 15 highest-connectivity candidates and had coding-partner sets showing significant GO and KEGG enrichment. Together, these findings prioritized the two lncRNAs through the combined evidence of differential expression, temporal responsiveness, connectivity, functional enrichment, and genomic proximity. These associations remain candidates for experimental investigation rather than established cis-regulatory relationships.

### Integrated coexpression network of temporally prioritized novel lncRNAs

Thirteen of the 14 temporally prioritized novel lncRNAs had significant coexpression associations, collectively forming 1,744 connections with 1,149 protein-coding genes. These included 896 positive and 848 negative associations. MSTRG.6052 had no coding gene partners meeting the coexpression criteria. Among the connected candidates, MSTRG.22879, MSTRG.6308, MSTRG.32011, and MSTRG.25627 had the highest connectivity, with 408, 335, 303, and 260 partners, respectively **(Supplementary Table S8)**. A reduced visualization retained the five highest-ranked coding-gene partners of each connected lncRNA together with all four prioritized cis-associated pairs **(Figure 5)**. The resulting network contained 13 novel lncRNAs, 64 protein-coding genes, and 66 associations **(Supplementary Table S9)**.

**Figure 5.**
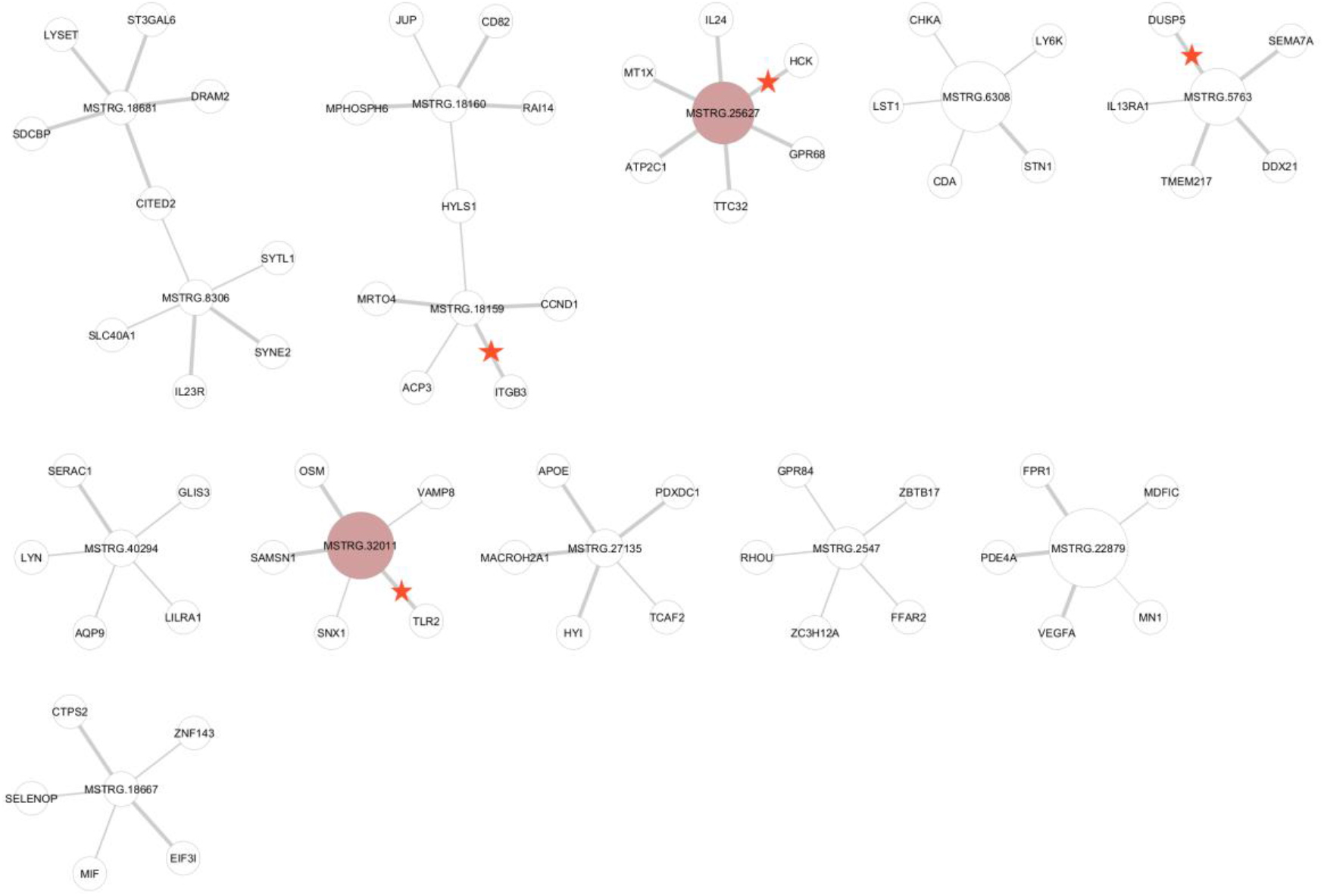
Coexpression network of temporally prioritized novel lncRNAs. The reduced network contains 13 novel lncRNAs, 64 protein-coding genes, and 66 associations (|r| ≥ 0.70; FDR < 0.05), retaining the five highest-ranked partners per lncRNA and all four supported cis-associated pairs. Pink nodes highlight MSTRG.32011 and MSTRG.25627; stars indicate cis-associated pairs. The node size reflects connectivity in the full network, and thicker edges indicate more positive correlations.

This representation combined temporal prioritization, coexpression, and genomic proximity while preserving the distinction between statistical associations and demonstrated regulatory interactions.

## Discussion

This study extends the original analysis of Candida auris-stimulated PBMCs by characterizing previously unannotated host lncRNA responses. The analysis identified 607 putative novel polyadenylated lncRNA loci, including 102 that were differentially expressed in at least one stimulus-time comparison. Among these genes, 94 were significantly coexpressed with protein-coding genes. Temporal interaction analysis identified 19 unique novel lncRNAs, 14 of which were also differentially expressed; therefore, this analysis was carried forward for integrated prioritization. Combining temporal responsiveness, coexpression, functional enrichment, and genomic proximity identified 4 candidate cis associations and highlighted MSTRG.32011 and MSTRG.25627 as the strongest candidates for experimental investigation.

The main novelty of this study is the identification and prioritization of human transcripts absent from GENCODE Release 48. Previous studies have examined host lncRNAs during *C. albicans* infection in mice ^9^ or lncRNAs encoded by C. auris itself ^11^. To our knowledge, this is the first systematic analysis of putative novel human lncRNAs in donor-matched PBMCs exposed to live C. auris and its purified cell wall components. Here, “novel” means absent from GENCODE Release 48 and does not imply absence from every transcript database or human tissue.

The transcriptional response increased substantially between 4 h and 24 h, with live C. auris producing the broadest late response **(Table 3)**. This finding is consistent with that of the original study, which revealed that β-glucan contributed strongly to the early response, whereas C. auris mannan was important during the later response ^4^. The novel lncRNAs showed a similar pattern, with most responses detected at 24 h. The 4 h and 24 h RPMI controls were separated according to the PCA, indicating a time-dependent shift in overall PBMC gene expression even without fungal stimulation. Therefore, each stimulated group was compared with its time-matched RPMI control group **(Supplementary Figure S1B).**

The novel-lncRNA response is frequently stimulus specific. At 24 h, 57 of the 78 lncRNAs that respond to live C. auris were unique to that contrast, whereas 16 of the 34 mannan-responsive lncRNAs were unique to mannan **(Figure 3B).** Only 10 were shared between the two contrasts. The broader response to live fungi may reflect the recognition of several cell wall components and additional signals generated during host-fungal interactions.

Coexpression analysis connected 94 of the 102 differentially expressed novel lncRNAs with protein-coding genes. The coding partners of the 15 most highly connected lncRNAs were enriched for defense, innate immune, and biotic-stimulus responses **(Figure 4B; Supplementary Table S5)**. Individual lncRNA partner sets were also associated with the lysosomal, NOD-like receptor, and JAK-STAT pathways. These findings are consistent with the involvement of lncRNAs in immune cell activation ^40^. However, the lack of significant combined KEGG or Reactome enrichment suggests that lncRNAs may participate in several distinct biological programs. Coexpression also indicates coordinated expression, not direct regulation.

Temporal interaction analysis revealed 19 lncRNAs whose responses changed between 4 h and 24 h, including 5 that were not identified as differentially expressed in any of the six stimulus versus RPMI contrasts analyzed separately at each time point **(Supplementary Table S6)**. Thus, temporal testing identified candidates that would have been missed by considering each time point separately.

The integrated network summarized these relationships among the temporally prioritized candidates. The reduced network contained 13 novel lncRNAs, 64 protein-coding genes, and 66 associations, including the four supported proximal pairs **(Figure 5)**. It therefore provides a focused map for selecting candidates for validation, but the edges represent statistical associations rather than confirmed molecular interactions.

The four proximal pairs presented differential expression, genomic proximity, and significant positive coexpression **(Table 4)**. MSTRG.32011 was located 3,922 bp from TLR2 and showed the strongest correlation (r = 0.937). This finding is biologically relevant because TLR2 is independently associated with the human transcriptional response to C. auris ^41^. MSTRG.25627 overlapped with the HCK gene-body interval and was also highly connected. HCK is a myeloid Src family kinase involved in phagocytosis, cell migration, actin organization, and lysosomal exocytosis ^42^. The other supported pairs were MSTRG.5763-DUSP5 and MSTRG.18159-ITGB3. Nevertheless, proximity and correlation cannot establish cis regulation. The effects of RNA molecules, transcriptional processes, or local regulatory DNA must be distinguished experimentally ^43^. The overlapping MSTRG.25627-HCK locus also requires confirmation of transcript boundaries.

The important **strengths** of this study include its balanced donor-matched design, consistently high sequencing quality, and multistage candidate-filtering strategy. Splice-junction validation, internal-priming and coding-potential filters, donor-supported expression, and manual genomic review reduced several common sources of false-positive lncRNA discovery. Quantification against a unified annotation also allowed novel and established genes to be evaluated within the same analysis.

Several **limitations** should be considered. Only 3 donors were included, and the 24 libraries do not represent 24 independent individuals. Therefore, the coexpression results should be used to prioritize candidate lncRNA-gene relationships rather than as evidence of direct biological regulation. In addition, 3′ QuantSeq is not suitable for resolving complete transcript structures or distant isoforms ^44^. The reported candidates should therefore be considered putative polyadenylated lncRNA loci rather than confirmed full-length transcripts. Bulk PBMC sequencing cannot determine which immune cell population expresses each lncRNA, and correlations may partly reflect differences among cell populations. Finally, the experiment included only two time points, one live C. auris strain, and PBMCs from healthy donors under ex vivo conditions. Immune responses may vary among cell types, fungal isolates, and clinical settings ^5^.

**Future studies** should validate candidate expression and strand orientation by strand-specific RT-PCR or RT-qPCR in a larger donor cohort. Transcript structures should be confirmed via full-length or long-read RNA sequencing. Additional time points, multiple C. auris strains or clades, and sorted or single-cell immune populations could clarify the specificity and cellular origin of these responses. Finally, CRISPR interference, antisense oligonucleotide, or related perturbation approaches are needed to determine whether prioritized lncRNAs influence nearby genes, immune pathways, cytokine production, or antifungal activity.

## Conclusion

This study identified a previously underexplored lncRNA component of the human response to C. auris. These responses are predominantly late, often stimulus specific, and associated with protein-coding programs involved in innate immunity. Although the results do not demonstrate regulatory function, they provide a focused set of candidates, particularly MSTRG.32011 and MSTRG.25627, for experimental investigation.

## Supplementary Information

Supplementary figures, tables, and supporting data are provided with this article.

## List of abbreviations

BH: Benjamini-Hochberg
BP: biological process
CPC2: Coding Potential Calculator 2
CPM: counts per million
DE: differentially expressed
FDR: false discovery rate
GO: Gene Ontology
IGV: Integrative Genomics Viewer
KEGG: Kyoto Encyclopedia of Genes and Genomes
lncRNA: long noncoding RNA
ORF: open reading frame
PBMC: peripheral blood mononuclear cell
PCA: principal component analysis
RNA-seq: RNA sequencing
RPMI: Roswell Park Memorial Institute medium
SRA: Sequence Read Archive
TMM: trimmed mean of M values

## Declarations

### Ethics approval and consent to participate

The present study involved secondary analysis of the publicly available dataset from NCBI BioProject PRJNA647871 and did not involve new participants or sample collection.

### Clinical trial number

Not applicable

### Consent for publication

Not applicable.

### Availability of data and materials

The RNA-sequencing data analyzed during the current study are publicly available through the NCBI Sequence Read Archive under BioProject accession number PRJNA647871 and through the Gene Expression Omnibus under accession number GSE154911. The analysis code, computational workflow, environment specifications, and supporting processed data have been archived in [Zenodo](https://zenodo.org/records/23028223). The current development version of the workflow is available through [GitHub](https://github.com/takim27/C.auris_transcriptomics)

### Authors’ contributions

**MTS** conceptualized the study, developed the methodology, performed the computational and statistical analyses, curated the data, generated the visualizations, and wrote the original manuscript. **BB** contributed to study conceptualization, supervised the research, interpreted the findings, and reviewed and edited the manuscript. Both authors read and approved the final manuscript.

## Acknowledgment

We thank Sean Harrington for the analytical assistance made possible by an Institutional Development Award (IDeA) from the National Institute of General Medical Sciences of the National Institutes of Health under Grant # 2P20GM103432.

## List of Supplementary Files

**Supplementary Data 1**. Interactive MultiQC HTML report summarizing FastQC results for the 24 untrimmed QuantSeq libraries, including read counts, read lengths, base-call quality, sequence composition, duplication and adapter content.

**Supplementary Data 2**. Interactive MultiQC HTML report summarizing quality-control results for the 24 libraries after Cutadapt processing, including retained read counts, read lengths, base-call quality and residual adapter content.

**Supplementary Data 3**. Interactive MultiQC HTML report summarizing STAR alignment metrics, including overall and unique mapping rates, multimapping, mismatch and insertion/deletion rates, and annotated and unannotated splice-junction counts.

**Supplementary Data 4**. Results of splice-junction validation, internal-priming assessment, Rfam screening, CPC2 coding-potential classification and Pfam domain screening. Candidate identifiers, screening results and pass or fail decisions are reported for each filtering stage.

**Supplementary Data 5**. Complete edgeR results for all genes tested in the six stimulus vs time matched RPMI comparisons: live C. auris, mannan and β-glucan at 4 and 24 h. Results include log₂ fold changes, expression levels, test statistics, P values, false-discovery rates and differential-expression classifications.

**Supplementary Figure S1**. (A) Library sizes for the 24 RNA-sequencing samples. (B) Principal-component analysis of normalized log₂-CPM values, with samples grouped by time, stimulus and donor. PC1 and PC2 explained 37.3% and 13.3% of the variance, respectively.

**Supplementary Figure S2**. UpSet plots showing intersections of upregulated and downregulated genes across the six stimulus-versus-time-matched RPMI comparisons, presented separately for all differentially expressed genes and novel lncRNAs.

**Supplementary Table S1**. Sample accessions, donor identities, stimulation conditions, time points and experimental-group assignments for the 24 RNA-sequencing libraries.

**Supplementary Table S2**. Genomic regions, candidate and coding-gene identifiers, spatial relationships, distances, coverage gaps, connecting junctions and manual retention decisions for the 34 transcript - coding gene relationships examined in IGV.

**Supplementary Table S3**. Distribution of GffCompare class codes in the merged transcriptome assembly, including the 388 intergenic transcripts assigned class code u and the 1,294 opposite-strand overlapping transcripts assigned class code x that formed the primary discovery set.

**Supplementary Table S4**. Maximum and mean CPM, maximum donor support, best-supported stimulus-time group and expression-screening outcome for each candidate locus. Candidates passed when CPM was ≥0.5 in at least two donors within the same experimental group.

**Supplementary Table S5**. Combined and individual-lncRNA enrichment results for Gene Ontology biological processes, KEGG pathways and Reactome pathways among the coding-gene partners of the 15 most highly connected novel lncRNAs.

**Supplementary Table S6**. Expression changes at 4 and 24 h and stimulus-by-time interaction results for 22 significant interactions involving 19 novel lncRNA loci. The table includes log₂ fold changes, FDR values, expression status, temporal-response classifications and prioritization indicators.

**Supplementary Table S7**. Genomic coordinates, strand relationships and gene-body distances for 60 proximal lncRNA-coding-gene pairs involving 13 novel lncRNAs and 56 distinct protein-coding genes.

**Supplementary Table S8**. Complete network containing 1,744 significant co-expression associations between 13 novel lncRNAs and 1,149 protein-coding genes, including 896 positive and 848 negative associations. Correlation statistics and genomic-proximity annotations are provided for each pair.

**Supplementary Table S9**. Reduced network containing the five strongest coding-gene partners of each connected lncRNA together with all four prioritized cis-associated pairs. The displayed network contains 13 novel lncRNAs, 64 protein-coding genes and 66 associations.

