## Supplementary material for "Computational identification of candidate human long noncoding RNAs associated with PBMC responses to *Candida auris* and its cell wall components": Supplementary_Data_2.html (from another version of this preprint: version 1, 2026-09-22)

Loading report..

Report
generated on 2026-07-29, 07:26 MDT
based on data in:
`/cluster/medbow/project/bishalab/msarker/c.auris_new/fastqc_trimmed`

Summarize report

Copy report prompt

**Welcome!** Not sure where to start?
Watch a tutorial video
*(6:06)*

don't show again

###### Report AI Summary

More details…

Provider: , model:

Chat with Seqera AI

### General Statistics

###### AI Summary

Provider: , model:

Chat with Seqera AI

Table
 Export...

Copy prompt


Summarize plot

Created with MultiQC

Copy table

 Configure columns

 Sort by highlight

 Scatter plot

 Violin plot
Export as CSV...
Showing ^24^/_24_ rows and ^3^/_6_ columns.

Copy Prompt


Summarize table

| Sample Name | Dups | GC | Avg len | Median len | Failed | Seqs |
| --- | --- | --- | --- | --- | --- | --- |
| SRR12291324 | 24.5% | 46.0% | 71bp | 74bp | 20% | 3.1M |
| SRR12291328 | 58.9% | 45.0% | 72bp | 74bp | 20% | 6.4M |
| SRR12291332 | 41.4% | 44.0% | 72bp | 74bp | 10% | 4.9M |
| SRR12291333 | 63.7% | 44.0% | 73bp | 74bp | 20% | 6.3M |
| SRR12291334 | 65.7% | 44.0% | 72bp | 74bp | 20% | 8.3M |
| SRR12291335 | 45.8% | 43.0% | 72bp | 74bp | 10% | 3.6M |
| SRR12291336 | 56.5% | 43.0% | 72bp | 74bp | 20% | 4.5M |
| SRR12291337 | 53.6% | 44.0% | 72bp | 74bp | 20% | 4.9M |
| SRR12291338 | 39.0% | 44.0% | 70bp | 74bp | 10% | 4.9M |
| SRR12291342 | 34.3% | 44.0% | 72bp | 74bp | 10% | 4.0M |
| SRR12291346 | 45.8% | 44.0% | 71bp | 74bp | 10% | 8.6M |
| SRR12291347 | 49.9% | 44.0% | 72bp | 74bp | 10% | 11.0M |
| SRR12291348 | 52.2% | 44.0% | 72bp | 74bp | 20% | 10.7M |
| SRR12291349 | 43.0% | 43.0% | 72bp | 74bp | 10% | 8.8M |
| SRR12291350 | 30.2% | 43.0% | 72bp | 74bp | 10% | 2.5M |
| SRR12291351 | 37.9% | 44.0% | 72bp | 74bp | 10% | 5.3M |
| SRR12291352 | 35.6% | 44.0% | 71bp | 74bp | 10% | 2.5M |
| SRR12291356 | 46.1% | 44.0% | 72bp | 74bp | 10% | 4.7M |
| SRR12291360 | 44.7% | 44.0% | 71bp | 74bp | 10% | 3.1M |
| SRR12291361 | 49.8% | 44.0% | 71bp | 74bp | 10% | 4.7M |
| SRR12291362 | 36.8% | 44.0% | 71bp | 74bp | 10% | 2.9M |
| SRR12291363 | 46.1% | 44.0% | 72bp | 74bp | 10% | 7.6M |
| SRR12291364 | 42.0% | 42.0% | 72bp | 74bp | 10% | 3.6M |
| SRR12291365 | 29.5% | 43.0% | 71bp | 74bp | 10% | 2.2M |

Copy Prompt


Summarize table

| Overrepresented sequence | Reports | Occurrences | % of all reads |
| --- | --- | --- | --- |
| CGCGACCTCAGATCAGACGTGGCGACCCGCTGAATTTAAGCATATTAGTC | 16 | 148904 | 0.1154% |
| GCGACCTCAGATCAGACGTGGCGACCCGCTGAATTTAAGCATATTAGTCA | 1 | 6241 | 0.0048% |

##### FastQC: Top overrepresented sequences: Columns

Uncheck the tick box to hide columns. Click and drag the handle on the left to change order. Table ID: `fastqc_top_overrepresented_sequences_table_table`

Show All
Show None

| Sort | Visible | Group | Column | Description | ID | Scale |
| --- | --- | --- | --- | --- | --- | --- |
| \|\| |  |  | Reports | Number of FastQC reports where this sequence is founds as overrepresented | `fastqc-samples` |  |
| \|\| |  |  | Occurrences | Total number of occurrences of the sequence (among the samples where the sequence is overrepresented) | `fastqc-total_count` |  |
| \|\| |  |  | % of all reads | Total number of occurrences as the percentage of all reads (among samples where the sequence is overrepresented) | `fastqc-total_percent` |  |

Close

#### Adapter Content 24 Help

The cumulative percentage count of the proportion of your
library which has seen each of the adapter sequences at each position.

**120 sample-adapter combinations** with less than 0.1% adapter contamination hidden from this plot across 24 samples.

Close
