## Supplementary material for "Computational identification of candidate human long noncoding RNAs associated with PBMC responses to *Candida auris* and its cell wall components": Supplementary_Data_3.html (from another version of this preprint: version 1, 2026-09-22)

MultiQC Report


Loading report..

v1.35


Theme

- Light
- Dark
- Auto

Highlight

 Rename

 Show / Hide


AI Analysis

 Export

 Settings

 Citations

 About

- General Stats
- STAR
  - Summary Statistics
  - Alignment Scores
  - Gene Counts

Toolbox

##### MultiQC Toolbox

Loading report..

Report
generated on 2026-07-29, 19:00 MDT
based on data in:
`/cluster/medbow/project/bishalab/msarker/c.auris_new/STAR_alignment`

Summarize report

Copy report prompt

**Welcome!** Not sure where to start?
Watch a tutorial video
*(6:06)*

don't show again

###### Report AI Summary

More details…

Provider: , model:

Chat with Seqera AI

### General Statistics

###### AI Summary

Provider: , model:

Chat with Seqera AI

Table
 Export...

Copy prompt


Summarize plot

Created with MultiQC

Copy table

 Configure columns

 Sort by highlight

 Scatter plot

 Violin plot
Export as CSV...
Showing ^48^/_48_ rows and ^2^/_6_ columns.

Copy Prompt


Summarize table

| Sample Name | Total reads | Aligned | Aligned | Uniq aligned | Uniq aligned | Multimapped |
| --- | --- | --- | --- | --- | --- | --- |
| SRR12291324 | 3.1M | 3.1M | 98.6% | 2.5M | 78.6% | 0.6M |
| SRR12291324\_\_STARpass1 | 3.1M | 3.1M | 98.6% | 2.4M | 78.6% | 0.6M |
| SRR12291328 | 6.4M | 6.4M | 99.4% | 4.9M | 75.9% | 1.5M |
| SRR12291328\_\_STARpass1 | 6.4M | 6.4M | 99.4% | 4.9M | 75.9% | 1.5M |
| SRR12291332 | 4.9M | 4.8M | 98.8% | 3.9M | 80.3% | 0.9M |
| SRR12291332\_\_STARpass1 | 4.9M | 4.8M | 98.8% | 3.9M | 80.4% | 0.9M |
| SRR12291333 | 6.3M | 6.2M | 99.3% | 5.0M | 79.8% | 1.2M |
| SRR12291333\_\_STARpass1 | 6.3M | 6.2M | 99.3% | 5.0M | 79.8% | 1.2M |
| SRR12291334 | 8.3M | 8.3M | 99.4% | 6.5M | 78.5% | 1.7M |
| SRR12291334\_\_STARpass1 | 8.3M | 8.3M | 99.4% | 6.5M | 78.6% | 1.7M |
| SRR12291335 | 3.6M | 3.5M | 99.3% | 2.9M | 82.5% | 0.6M |
| SRR12291335\_\_STARpass1 | 3.6M | 3.5M | 99.3% | 2.9M | 82.5% | 0.6M |
| SRR12291336 | 4.5M | 4.5M | 99.4% | 3.7M | 82.5% | 0.8M |
| SRR12291336\_\_STARpass1 | 4.5M | 4.5M | 99.4% | 3.7M | 82.5% | 0.8M |
| SRR12291337 | 4.9M | 4.8M | 98.9% | 4.0M | 81.5% | 0.8M |
| SRR12291337\_\_STARpass1 | 4.9M | 4.8M | 98.9% | 4.0M | 81.5% | 0.8M |
| SRR12291338 | 4.9M | 4.9M | 99.3% | 3.8M | 78.2% | 1.0M |
| SRR12291338\_\_STARpass1 | 4.9M | 4.9M | 99.3% | 3.8M | 78.2% | 1.0M |
| SRR12291342 | 4.0M | 4.0M | 99.3% | 3.3M | 80.9% | 0.7M |
| SRR12291342\_\_STARpass1 | 4.0M | 4.0M | 99.3% | 3.3M | 81.0% | 0.7M |
| SRR12291346 | 8.6M | 8.5M | 99.4% | 6.8M | 79.6% | 1.7M |
| SRR12291346\_\_STARpass1 | 8.6M | 8.5M | 99.4% | 6.9M | 79.8% | 1.7M |
| SRR12291347 | 11.0M | 10.9M | 99.3% | 8.9M | 81.1% | 2.0M |
| SRR12291347\_\_STARpass1 | 11.0M | 10.9M | 99.3% | 8.9M | 81.2% | 2.0M |
| SRR12291348 | 10.7M | 10.6M | 99.4% | 8.5M | 79.5% | 2.1M |
| SRR12291348\_\_STARpass1 | 10.7M | 10.6M | 99.4% | 8.5M | 79.7% | 2.1M |
| SRR12291349 | 8.8M | 8.7M | 99.1% | 7.3M | 82.8% | 1.4M |
| SRR12291349\_\_STARpass1 | 8.8M | 8.7M | 99.1% | 7.3M | 82.9% | 1.4M |
| SRR12291350 | 2.5M | 2.5M | 99.4% | 2.0M | 82.3% | 0.4M |
| SRR12291350\_\_STARpass1 | 2.5M | 2.5M | 99.4% | 2.0M | 82.4% | 0.4M |
| SRR12291351 | 5.3M | 5.3M | 99.3% | 4.3M | 81.8% | 0.9M |
| SRR12291351\_\_STARpass1 | 5.3M | 5.3M | 99.3% | 4.3M | 81.8% | 0.9M |
| SRR12291352 | 2.5M | 2.5M | 99.4% | 2.0M | 79.6% | 0.5M |
| SRR12291352\_\_STARpass1 | 2.5M | 2.5M | 99.4% | 2.0M | 79.8% | 0.5M |
| SRR12291356 | 4.7M | 4.7M | 99.2% | 3.9M | 82.3% | 0.8M |
| SRR12291356\_\_STARpass1 | 4.7M | 4.7M | 99.2% | 3.9M | 82.3% | 0.8M |
| SRR12291360 | 3.1M | 3.1M | 99.1% | 2.4M | 78.4% | 0.6M |
| SRR12291360\_\_STARpass1 | 3.1M | 3.1M | 99.1% | 2.4M | 78.5% | 0.6M |
| SRR12291361 | 4.7M | 4.7M | 99.5% | 3.6M | 77.0% | 1.1M |
| SRR12291361\_\_STARpass1 | 4.7M | 4.7M | 99.4% | 3.6M | 77.2% | 1.1M |
| SRR12291362 | 2.9M | 2.9M | 99.3% | 2.3M | 79.1% | 0.6M |
| SRR12291362\_\_STARpass1 | 2.9M | 2.9M | 99.3% | 2.3M | 79.3% | 0.6M |
| SRR12291363 | 7.6M | 7.5M | 98.9% | 6.1M | 80.5% | 1.4M |
| SRR12291363\_\_STARpass1 | 7.6M | 7.5M | 98.9% | 6.1M | 80.6% | 1.4M |
| SRR12291364 | 3.6M | 3.6M | 99.1% | 3.1M | 84.0% | 0.6M |
| SRR12291364\_\_STARpass1 | 3.6M | 3.6M | 99.1% | 3.1M | 84.0% | 0.5M |
| SRR12291365 | 2.2M | 2.1M | 99.3% | 1.8M | 83.1% | 0.3M |
| SRR12291365\_\_STARpass1 | 2.2M | 2.1M | 99.3% | 1.8M | 83.1% | 0.3M |

Expand table

##### General Statistics: Columns

Uncheck the tick box to hide columns. Click and drag the handle on the left to change order. Table ID: `general_stats_table_table`

Show All
Show None

| Sort | Visible | Group | Column | Description | ID | Scale |
| --- | --- | --- | --- | --- | --- | --- |
| \|\| |  | STAR | Total reads | Number of input reads | `star-total_reads` | read\_count |
| \|\| |  | STAR | Aligned | Mapped reads | `star-mapped` | read\_count |
| \|\| |  | STAR | Aligned | % Mapped reads | `star-mapped_percent` |  |
| \|\| |  | STAR | Uniq aligned | Uniquely mapped reads | `star-uniquely_mapped` | read\_count |
| \|\| |  | STAR | Uniq aligned | % Uniquely mapped reads | `star-uniquely_mapped_percent` |  |
| \|\| |  | STAR | Multimapped | Multiple mapped reads | `star-multimapped` | read\_count |

Close

### STAR

Universal RNA-seq aligner.https://github.com/alexdobin/STARDOI: 10.1093/bioinformatics/bts635

#### Summary Statistics

Summary statistics from the STAR alignment

###### AI Summary

Provider: , model:

Chat with Seqera AI

Table
 Export...

Copy prompt


Summarize plot

Created with MultiQC

Copy table

 Configure columns

 Sort by highlight

 Scatter plot

 Violin plot
Export as CSV...
Showing ^48^/_48_ rows and ^10^/_19_ columns.

Copy Prompt


Summarize table

| Sample Name | Total reads | Aligned | Aligned | Uniq aligned | Uniq aligned | Multimapped | Avg. read len | Avg. mapped len | Splices | Annotated splices | GT/AG splices | GC/AG splices | AT/AC splices | Non-canonical splices | Mismatch rate | Del rate | Del len | Ins rate | Ins len |
| --- | --- | --- | --- | --- | --- | --- | --- | --- | --- | --- | --- | --- | --- | --- | --- | --- | --- | --- | --- |
| SRR12291324 | 3.1M | 3.1M | 98.6% | 2.5M | 78.6% | 0.6M | 71.0bp | 70.3bp | 0.1M | 0.1M | 0.1M | 0.0M | 0.0M | 0.0M | 0.5% | 0.0% | 1.4bp | 0.0% | 1.4bp |
| SRR12291324\_\_STARpass1 | 3.1M | 3.1M | 98.6% | 2.4M | 78.6% | 0.6M | 71.0bp | 70.3bp | 0.1M | 0.1M | 0.1M | 0.0M | 0.0M | 0.0M | 0.5% | 0.0% | 1.4bp | 0.0% | 1.4bp |
| SRR12291328 | 6.4M | 6.4M | 99.4% | 4.9M | 75.9% | 1.5M | 72.0bp | 71.2bp | 0.5M | 0.5M | 0.5M | 0.0M | 0.0M | 0.0M | 0.5% | 0.0% | 1.4bp | 0.0% | 1.3bp |
| SRR12291328\_\_STARpass1 | 6.4M | 6.4M | 99.4% | 4.9M | 75.9% | 1.5M | 72.0bp | 71.2bp | 0.5M | 0.5M | 0.5M | 0.0M | 0.0M | 0.0M | 0.5% | 0.0% | 1.4bp | 0.0% | 1.3bp |
| SRR12291332 | 4.9M | 4.8M | 98.8% | 3.9M | 80.3% | 0.9M | 71.0bp | 70.8bp | 0.4M | 0.4M | 0.4M | 0.0M | 0.0M | 0.0M | 0.6% | 0.0% | 1.5bp | 0.0% | 1.4bp |
| SRR12291332\_\_STARpass1 | 4.9M | 4.8M | 98.8% | 3.9M | 80.4% | 0.9M | 71.0bp | 70.8bp | 0.4M | 0.4M | 0.4M | 0.0M | 0.0M | 0.0M | 0.6% | 0.0% | 1.5bp | 0.0% | 1.4bp |
| SRR12291333 | 6.3M | 6.2M | 99.3% | 5.0M | 79.8% | 1.2M | 72.0bp | 71.4bp | 0.6M | 0.6M | 0.6M | 0.0M | 0.0M | 0.0M | 0.5% | 0.0% | 1.5bp | 0.0% | 1.4bp |
| SRR12291333\_\_STARpass1 | 6.3M | 6.2M | 99.3% | 5.0M | 79.8% | 1.2M | 72.0bp | 71.4bp | 0.6M | 0.6M | 0.6M | 0.0M | 0.0M | 0.0M | 0.5% | 0.0% | 1.5bp | 0.0% | 1.4bp |
| SRR12291334 | 8.3M | 8.3M | 99.4% | 6.5M | 78.5% | 1.7M | 72.0bp | 71.3bp | 0.9M | 0.8M | 0.8M | 0.0M | 0.0M | 0.0M | 0.5% | 0.0% | 1.5bp | 0.0% | 1.4bp |
| SRR12291334\_\_STARpass1 | 8.3M | 8.3M | 99.4% | 6.5M | 78.6% | 1.7M | 72.0bp | 71.3bp | 0.8M | 0.8M | 0.8M | 0.0M | 0.0M | 0.0M | 0.5% | 0.0% | 1.5bp | 0.0% | 1.4bp |
| SRR12291335 | 3.6M | 3.5M | 99.3% | 2.9M | 82.5% | 0.6M | 71.0bp | 70.5bp | 0.3M | 0.3M | 0.3M | 0.0M | 0.0M | 0.0M | 0.6% | 0.0% | 1.4bp | 0.0% | 1.3bp |
| SRR12291335\_\_STARpass1 | 3.6M | 3.5M | 99.3% | 2.9M | 82.5% | 0.6M | 71.0bp | 70.5bp | 0.3M | 0.3M | 0.3M | 0.0M | 0.0M | 0.0M | 0.6% | 0.0% | 1.4bp | 0.0% | 1.3bp |
| SRR12291336 | 4.5M | 4.5M | 99.4% | 3.7M | 82.5% | 0.8M | 72.0bp | 70.8bp | 0.3M | 0.3M | 0.3M | 0.0M | 0.0M | 0.0M | 0.5% | 0.0% | 1.4bp | 0.0% | 1.4bp |
| SRR12291336\_\_STARpass1 | 4.5M | 4.5M | 99.4% | 3.7M | 82.5% | 0.8M | 72.0bp | 70.8bp | 0.3M | 0.3M | 0.3M | 0.0M | 0.0M | 0.0M | 0.5% | 0.0% | 1.4bp | 0.0% | 1.4bp |
| SRR12291337 | 4.9M | 4.8M | 98.9% | 4.0M | 81.5% | 0.8M | 72.0bp | 70.9bp | 0.4M | 0.4M | 0.4M | 0.0M | 0.0M | 0.0M | 0.6% | 0.0% | 1.5bp | 0.0% | 1.3bp |
| SRR12291337\_\_STARpass1 | 4.9M | 4.8M | 98.9% | 4.0M | 81.5% | 0.8M | 72.0bp | 70.9bp | 0.4M | 0.4M | 0.4M | 0.0M | 0.0M | 0.0M | 0.6% | 0.0% | 1.5bp | 0.0% | 1.3bp |
| SRR12291338 | 4.9M | 4.9M | 99.3% | 3.8M | 78.2% | 1.0M | 70.0bp | 69.6bp | 0.4M | 0.4M | 0.4M | 0.0M | 0.0M | 0.0M | 0.5% | 0.0% | 1.4bp | 0.0% | 1.3bp |
| SRR12291338\_\_STARpass1 | 4.9M | 4.9M | 99.3% | 3.8M | 78.2% | 1.0M | 70.0bp | 69.6bp | 0.4M | 0.4M | 0.4M | 0.0M | 0.0M | 0.0M | 0.5% | 0.0% | 1.4bp | 0.0% | 1.3bp |
| SRR12291342 | 4.0M | 4.0M | 99.3% | 3.3M | 80.9% | 0.7M | 72.0bp | 71.1bp | 0.3M | 0.3M | 0.3M | 0.0M | 0.0M | 0.0M | 0.5% | 0.0% | 1.4bp | 0.0% | 1.3bp |
| SRR12291342\_\_STARpass1 | 4.0M | 4.0M | 99.3% | 3.3M | 81.0% | 0.7M | 72.0bp | 71.1bp | 0.3M | 0.3M | 0.3M | 0.0M | 0.0M | 0.0M | 0.5% | 0.0% | 1.4bp | 0.0% | 1.3bp |
| SRR12291346 | 8.6M | 8.5M | 99.4% | 6.8M | 79.6% | 1.7M | 71.0bp | 70.2bp | 0.7M | 0.7M | 0.7M | 0.0M | 0.0M | 0.0M | 0.5% | 0.0% | 1.4bp | 0.0% | 1.3bp |
| SRR12291346\_\_STARpass1 | 8.6M | 8.5M | 99.4% | 6.9M | 79.8% | 1.7M | 71.0bp | 70.2bp | 0.7M | 0.7M | 0.7M | 0.0M | 0.0M | 0.0M | 0.5% | 0.0% | 1.4bp | 0.0% | 1.3bp |
| SRR12291347 | 11.0M | 10.9M | 99.3% | 8.9M | 81.1% | 2.0M | 71.0bp | 70.6bp | 0.9M | 0.9M | 0.9M | 0.0M | 0.0M | 0.0M | 0.5% | 0.0% | 1.4bp | 0.0% | 1.3bp |
| SRR12291347\_\_STARpass1 | 11.0M | 10.9M | 99.3% | 8.9M | 81.2% | 2.0M | 71.0bp | 70.5bp | 0.9M | 0.9M | 0.9M | 0.0M | 0.0M | 0.0M | 0.5% | 0.0% | 1.4bp | 0.0% | 1.3bp |
| SRR12291348 | 10.7M | 10.6M | 99.4% | 8.5M | 79.5% | 2.1M | 71.0bp | 70.7bp | 1.0M | 1.0M | 1.0M | 0.0M | 0.0M | 0.0M | 0.5% | 0.0% | 1.4bp | 0.0% | 1.3bp |
| SRR12291348\_\_STARpass1 | 10.7M | 10.6M | 99.4% | 8.5M | 79.7% | 2.1M | 71.0bp | 70.7bp | 1.0M | 0.9M | 1.0M | 0.0M | 0.0M | 0.0M | 0.5% | 0.0% | 1.4bp | 0.0% | 1.3bp |
| SRR12291349 | 8.8M | 8.7M | 99.1% | 7.3M | 82.8% | 1.4M | 71.0bp | 70.5bp | 0.6M | 0.6M | 0.6M | 0.0M | 0.0M | 0.0M | 0.5% | 0.0% | 1.4bp | 0.0% | 1.3bp |
| SRR12291349\_\_STARpass1 | 8.8M | 8.7M | 99.1% | 7.3M | 82.9% | 1.4M | 71.0bp | 70.5bp | 0.6M | 0.6M | 0.6M | 0.0M | 0.0M | 0.0M | 0.5% | 0.0% | 1.4bp | 0.0% | 1.3bp |
| SRR12291350 | 2.5M | 2.5M | 99.4% | 2.0M | 82.3% | 0.4M | 71.0bp | 70.6bp | 0.2M | 0.2M | 0.2M | 0.0M | 0.0M | 0.0M | 0.5% | 0.0% | 1.4bp | 0.0% | 1.3bp |
| SRR12291350\_\_STARpass1 | 2.5M | 2.5M | 99.4% | 2.0M | 82.4% | 0.4M | 71.0bp | 70.6bp | 0.2M | 0.2M | 0.2M | 0.0M | 0.0M | 0.0M | 0.5% | 0.0% | 1.4bp | 0.0% | 1.3bp |
| SRR12291351 | 5.3M | 5.3M | 99.3% | 4.3M | 81.8% | 0.9M | 71.0bp | 70.7bp | 0.4M | 0.4M | 0.4M | 0.0M | 0.0M | 0.0M | 0.5% | 0.0% | 1.4bp | 0.0% | 1.3bp |
| SRR12291351\_\_STARpass1 | 5.3M | 5.3M | 99.3% | 4.3M | 81.8% | 0.9M | 71.0bp | 70.7bp | 0.4M | 0.4M | 0.4M | 0.0M | 0.0M | 0.0M | 0.5% | 0.0% | 1.4bp | 0.0% | 1.3bp |
| SRR12291352 | 2.5M | 2.5M | 99.4% | 2.0M | 79.6% | 0.5M | 71.0bp | 70.3bp | 0.2M | 0.2M | 0.2M | 0.0M | 0.0M | 0.0M | 0.6% | 0.0% | 1.4bp | 0.0% | 1.3bp |
| SRR12291352\_\_STARpass1 | 2.5M | 2.5M | 99.4% | 2.0M | 79.8% | 0.5M | 71.0bp | 70.3bp | 0.2M | 0.2M | 0.2M | 0.0M | 0.0M | 0.0M | 0.6% | 0.0% | 1.4bp | 0.0% | 1.3bp |
| SRR12291356 | 4.7M | 4.7M | 99.2% | 3.9M | 82.3% | 0.8M | 71.0bp | 70.7bp | 0.4M | 0.4M | 0.3M | 0.0M | 0.0M | 0.0M | 0.6% | 0.0% | 1.4bp | 0.0% | 1.3bp |
| SRR12291356\_\_STARpass1 | 4.7M | 4.7M | 99.2% | 3.9M | 82.3% | 0.8M | 71.0bp | 70.7bp | 0.3M | 0.3M | 0.3M | 0.0M | 0.0M | 0.0M | 0.6% | 0.0% | 1.4bp | 0.0% | 1.3bp |
| SRR12291360 | 3.1M | 3.1M | 99.1% | 2.4M | 78.4% | 0.6M | 71.0bp | 70.4bp | 0.3M | 0.3M | 0.3M | 0.0M | 0.0M | 0.0M | 0.6% | 0.0% | 1.4bp | 0.0% | 1.3bp |
| SRR12291360\_\_STARpass1 | 3.1M | 3.1M | 99.1% | 2.4M | 78.5% | 0.6M | 71.0bp | 70.4bp | 0.3M | 0.3M | 0.3M | 0.0M | 0.0M | 0.0M | 0.6% | 0.0% | 1.4bp | 0.0% | 1.3bp |
| SRR12291361 | 4.7M | 4.7M | 99.5% | 3.6M | 77.0% | 1.1M | 71.0bp | 70.3bp | 0.4M | 0.4M | 0.4M | 0.0M | 0.0M | 0.0M | 0.5% | 0.0% | 1.4bp | 0.0% | 1.3bp |
| SRR12291361\_\_STARpass1 | 4.7M | 4.7M | 99.4% | 3.6M | 77.2% | 1.1M | 71.0bp | 70.3bp | 0.4M | 0.4M | 0.4M | 0.0M | 0.0M | 0.0M | 0.5% | 0.0% | 1.4bp | 0.0% | 1.3bp |
| SRR12291362 | 2.9M | 2.9M | 99.3% | 2.3M | 79.1% | 0.6M | 71.0bp | 70.3bp | 0.2M | 0.2M | 0.2M | 0.0M | 0.0M | 0.0M | 0.5% | 0.0% | 1.4bp | 0.0% | 1.3bp |
| SRR12291362\_\_STARpass1 | 2.9M | 2.9M | 99.3% | 2.3M | 79.3% | 0.6M | 71.0bp | 70.3bp | 0.2M | 0.2M | 0.2M | 0.0M | 0.0M | 0.0M | 0.5% | 0.0% | 1.4bp | 0.0% | 1.3bp |
| SRR12291363 | 7.6M | 7.5M | 98.9% | 6.1M | 80.5% | 1.4M | 71.0bp | 70.7bp | 0.6M | 0.6M | 0.6M | 0.0M | 0.0M | 0.0M | 0.6% | 0.0% | 1.4bp | 0.0% | 1.3bp |
| SRR12291363\_\_STARpass1 | 7.6M | 7.5M | 98.9% | 6.1M | 80.6% | 1.4M | 71.0bp | 70.7bp | 0.6M | 0.6M | 0.6M | 0.0M | 0.0M | 0.0M | 0.6% | 0.0% | 1.4bp | 0.0% | 1.3bp |
| SRR12291364 | 3.6M | 3.6M | 99.1% | 3.1M | 84.0% | 0.6M | 71.0bp | 70.7bp | 0.3M | 0.3M | 0.3M | 0.0M | 0.0M | 0.0M | 0.6% | 0.0% | 1.4bp | 0.0% | 1.3bp |
| SRR12291364\_\_STARpass1 | 3.6M | 3.6M | 99.1% | 3.1M | 84.0% | 0.5M | 71.0bp | 70.7bp | 0.3M | 0.3M | 0.3M | 0.0M | 0.0M | 0.0M | 0.6% | 0.0% | 1.4bp | 0.0% | 1.3bp |
| SRR12291365 | 2.2M | 2.1M | 99.3% | 1.8M | 83.1% | 0.3M | 71.0bp | 70.4bp | 0.2M | 0.2M | 0.2M | 0.0M | 0.0M | 0.0M | 0.6% | 0.0% | 1.4bp | 0.0% | 1.3bp |
| SRR12291365\_\_STARpass1 | 2.2M | 2.1M | 99.3% | 1.8M | 83.1% | 0.3M | 71.0bp | 70.4bp | 0.2M | 0.1M | 0.1M | 0.0M | 0.0M | 0.0M | 0.6% | 0.0% | 1.4bp | 0.0% | 1.3bp |

Expand table

##### STAR: Summary Statistics: Columns

Uncheck the tick box to hide columns. Click and drag the handle on the left to change order. Table ID: `star_summary_table_table`

Show All
Show None

| Sort | Visible | Group | Column | Description | ID | Scale |
| --- | --- | --- | --- | --- | --- | --- |
| \|\| |  |  | Total reads | Number of input reads | `star-total_reads` | read\_count |
| \|\| |  |  | Aligned | Mapped reads | `star-mapped` | read\_count |
| \|\| |  |  | Aligned | % Mapped reads | `star-mapped_percent` |  |
| \|\| |  |  | Uniq aligned | Uniquely mapped reads | `star-uniquely_mapped` | read\_count |
| \|\| |  |  | Uniq aligned | % Uniquely mapped reads | `star-uniquely_mapped_percent` |  |
| \|\| |  |  | Multimapped | Multiple mapped reads | `star-multimapped` | read\_count |
| \|\| |  |  | Avg. read len | Average input read length | `star-avg_input_read_length` |  |
| \|\| |  |  | Avg. mapped len | Average mapped length | `star-avg_mapped_read_length` |  |
| \|\| |  |  | Splices | Number of splices: Total | `star-num_splices` | read\_count |
| \|\| |  |  | Annotated splices | Number of splices: Annotated (sjdb) | `star-num_annotated_splices` | read\_count |
| \|\| |  |  | GT/AG splices | Number of splices: GT/AG | `star-num_GTAG_splices` | read\_count |
| \|\| |  |  | GC/AG splices | Number of splices: GC/AG | `star-num_GCAG_splices` | read\_count |
| \|\| |  |  | AT/AC splices | Number of splices: AT/AC | `star-num_ATAC_splices` | read\_count |
| \|\| |  |  | Non-canonical splices | Number of splices: Non-canonical | `star-num_noncanonical_splices` | read\_count |
| \|\| |  |  | Mismatch rate | Mismatch rate per base | `star-mismatch_rate` |  |
| \|\| |  |  | Del rate | Deletion rate per base | `star-deletion_rate` |  |
| \|\| |  |  | Del len | Deletion average length | `star-deletion_length` |  |
| \|\| |  |  | Ins rate | Insertion rate per base | `star-insertion_rate` |  |
| \|\| |  |  | Ins len | Insertion average length | `star-insertion_length` |  |

Close

#### Alignment Scores

###### AI Summary

Provider: , model:

Chat with Seqera AI

Percentages
 Export...

Copy prompt


Summarize plot

Created with MultiQC

#### Gene Counts

Statistics from results generated using `--quantMode GeneCounts`. The three tabs show counts for unstranded RNA-seq, counts for the 1st read strand aligned with RNA and counts for the 2nd read strand aligned with RNA.

###### AI Summary

Provider: , model:

Chat with Seqera AI

Percentages

Unstranded
Same Stranded
Reverse Stranded

 Export...

Copy prompt


Summarize plot

Created with MultiQC

**MultiQC v1.35** 
- Written by Phil Ewels, available on
GitHub.

This report uses Plotly,
jQuery,
jQuery UI,
Bootstrap and
FileSaver.js.

#### Plot Table Data

Select Column

Select Column

Please select two table columns.

Close

#### Regex Help

Toolbox search strings can behave as regular expressions (regexes). Click a button below to see an example of
it in action. Try modifying them yourself in the text box.

`^` (start of string)

`$` (end of string)

`[]` (character choice)

`\d` (shorthand for `[0-9]`)

`\w` (shorthand for `[0-9a-zA-Z_]`)
`.` (any character)

`\.` (literal full stop)

`()` `|` (group / separator)

`*` (prev char 0 or more)

`+` (prev char 1 or more)

`?` (prev char 0 or 1)

`{}` (char num times)

`{,}` (count range)

```
samp_1
samp_1_edited
samp_2
samp_2_edited
samp_3
samp_3_edited
prepended_samp_1
tmp_samp_1_edited
tmpp_samp_1_edited
tmppp_samp_1_edited
#samp_1_edited.tmp
samp_11
samp_11111
```

See regex101.com for a more heavy duty testing suite.

Close
