## Supplementary material for "Computational identification of candidate human long noncoding RNAs associated with PBMC responses to *Candida auris* and its cell wall components": Supplementary_Figure_S1.docx (from another version of this preprint: version 1, 2026-09-22)

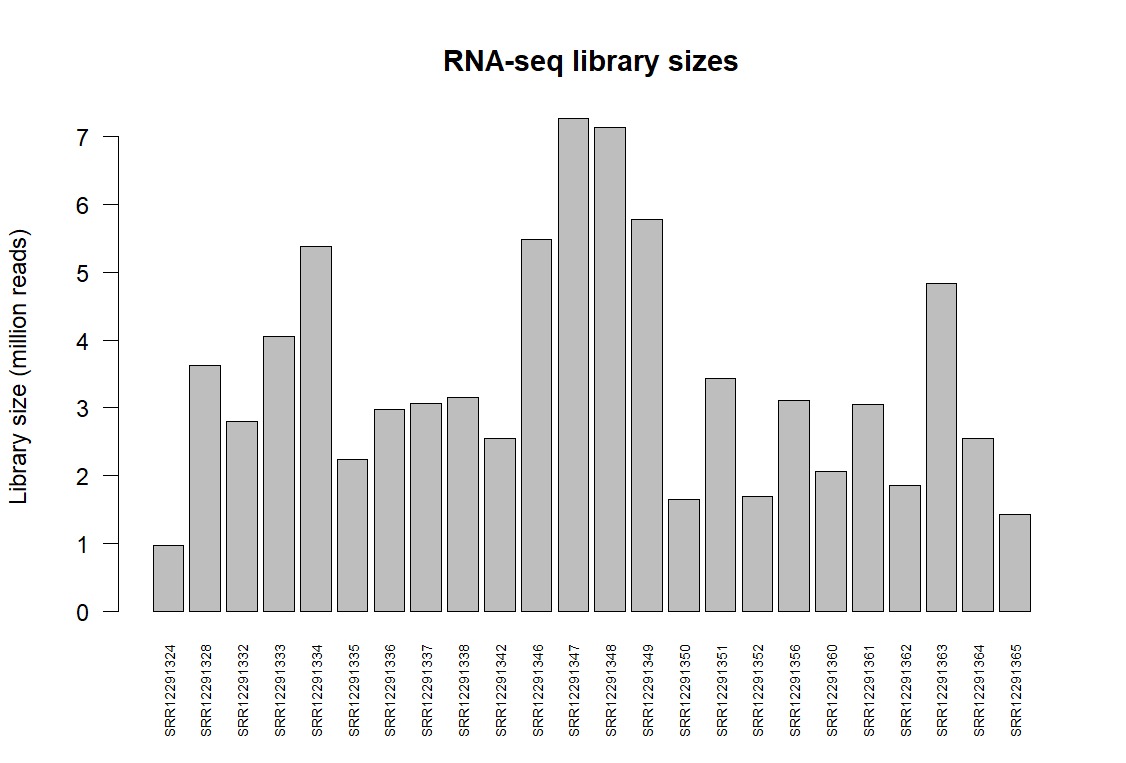

**Supplementary Figure S1(A). RNA-seq library sizes across the 24 samples.** Bars show the total reads assigned to expression-filtered genes, expressed in millions.

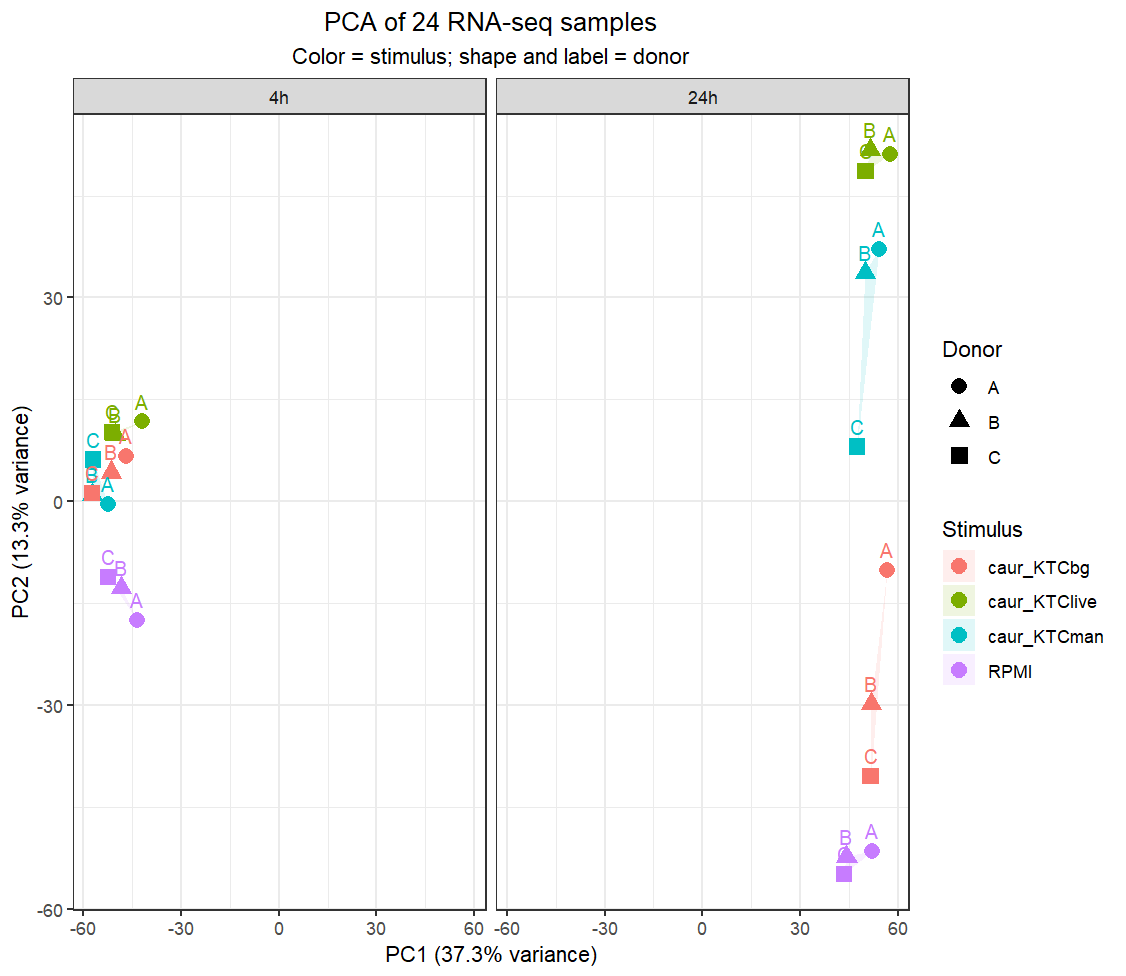

**Supplementary Figure S1(B). Principal-component analysis of the 24 RNA-seq libraries.** PCA was performed using normalized log₂-CPM values for 13,321 expression-filtered genes. Panels show 4 h and 24 h samples from the same PCA. Colors indicate β-glucan (coral), live C. auris (green), mannan (cyan), and RPMI (purple); shapes and labels identify donors A-C. Shaded polygons enclose samples within each stimulus-time group. PC1 and PC2 explain 37.3% and 13.3% of the variance, respectively.
