## Supplementary material for "Computational identification of candidate human long noncoding RNAs associated with PBMC responses to *Candida auris* and its cell wall components": Supplementary_Figure_S2.docx (from another version of this preprint: version 1, 2026-09-22)

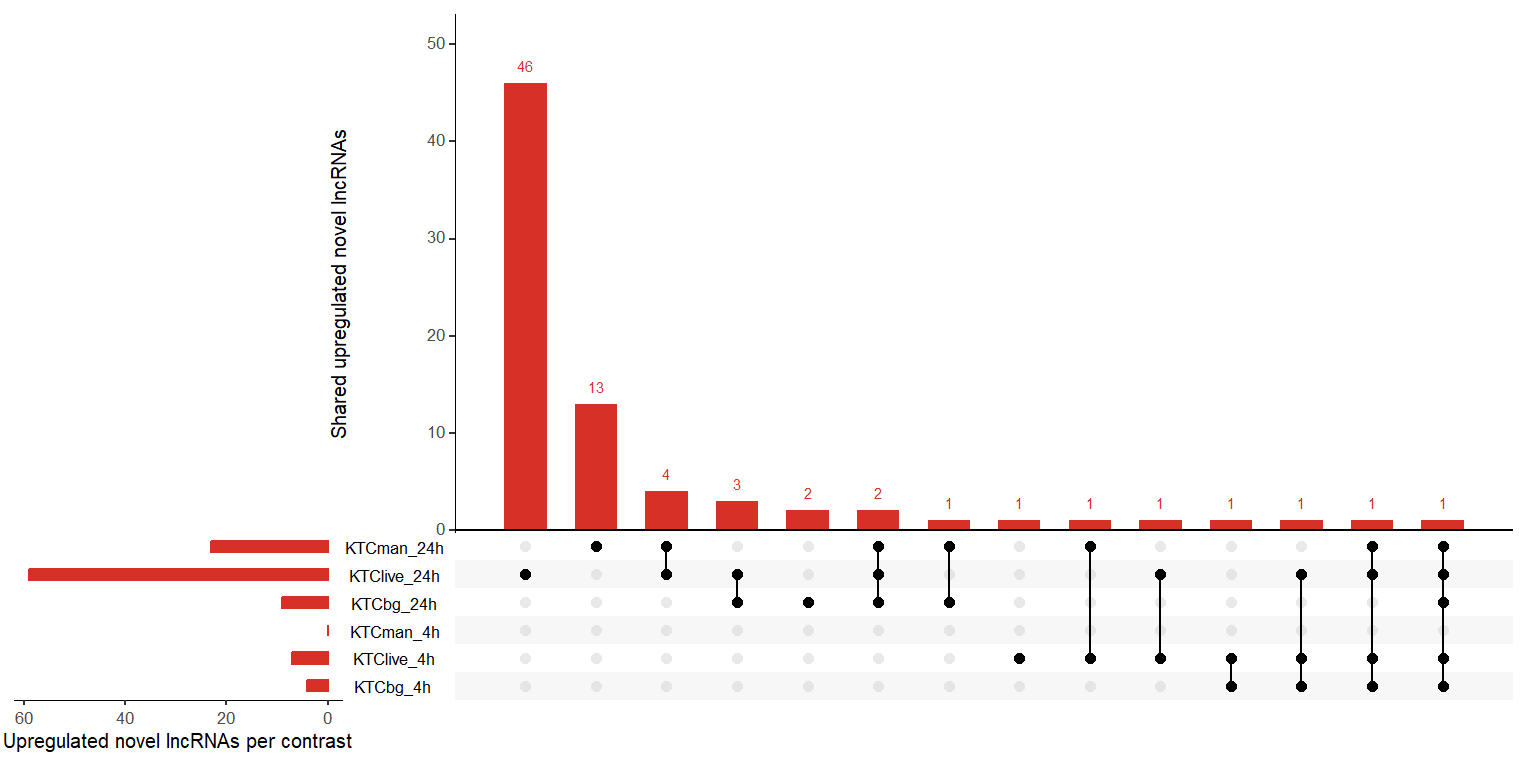

A

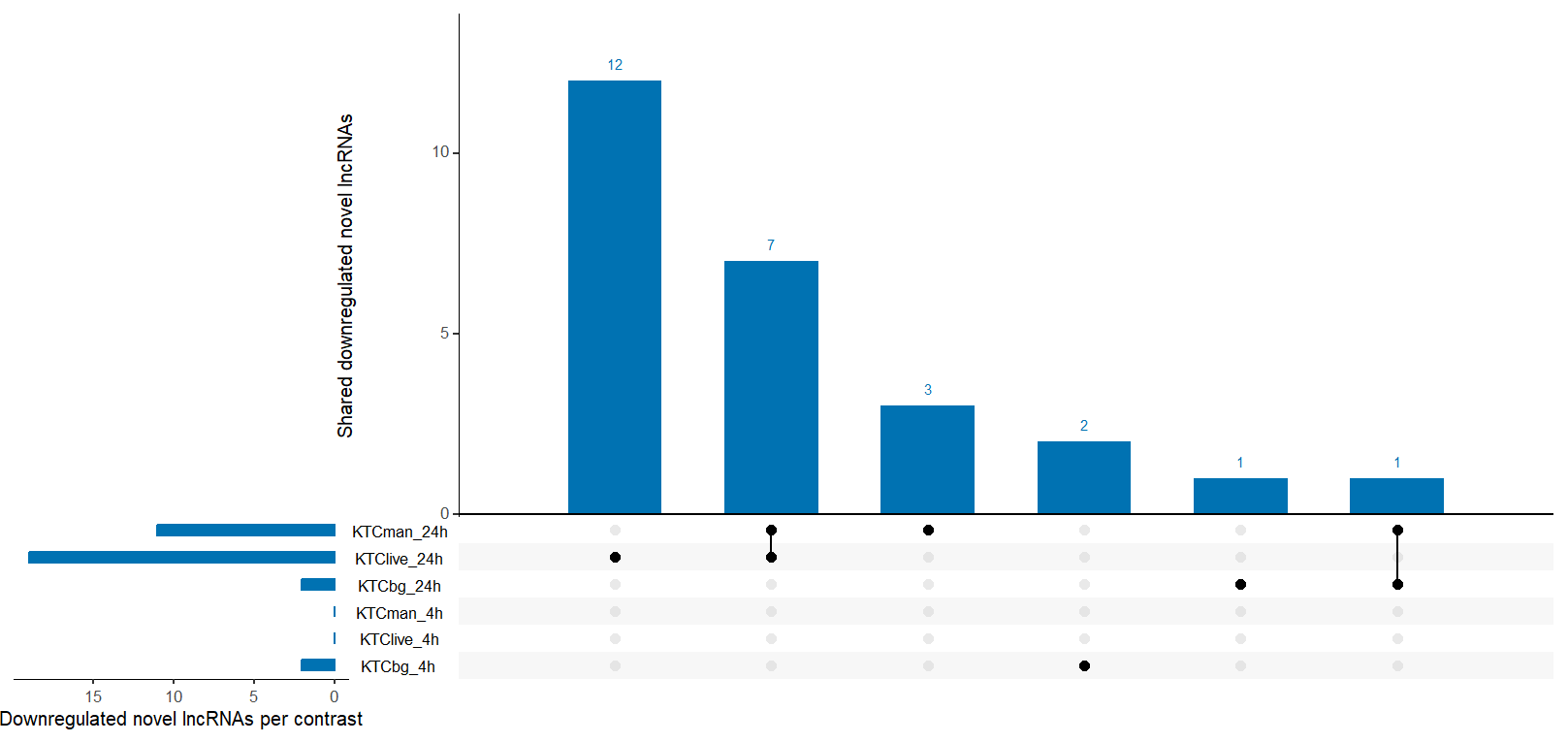

B

**Supplementary Figure S2.** Direction-specific overlap of differentially expressed novel lncRNAs across stimulus-time contrasts. UpSet plots show intersections among **(A)** upregulated and **(B)** downregulated novel lncRNAs following β-glucan (KTCbg), live *Candida auris* (KTClive), or mannan (KTCman) stimulation relative to the corresponding time-matched RPMI controls at 4 and 24 h. Horizontal bars indicate the total number of differentially expressed novel lncRNAs in each contrast, vertical bars indicate intersection sizes, and connected filled circles identify the contrasts contributing to each intersection. Differential expression was defined as FDR < 0.05 and |log2FC| ≥ 1.
