## Supplementary material for "Computational identification of candidate human long noncoding RNAs associated with PBMC responses to *Candida auris* and its cell wall components": Supplementary_Table_S1.docx (from another version of this preprint: version 1, 2026-09-22)

| Sample Name | Donor | GEO_Accession (exp) | stimulus | Type | time |
| --- | --- | --- | --- | --- | --- |
| SRR12291324 | Donor A | GSM4683203 | A1_RPMI_4hrs_100ng | Control | 4hrs |
| SRR12291328 | Donor A | GSM4683207 | A5_RPMI_24hrs_100ng | Control | 24hrs |
| SRR12291332 | Donor A | GSM4683211 | A9_caur_KTClive_4hrs_100ng | Test | 4hrs |
| SRR12291333 | Donor A | GSM4683212 | A10_caur_KTCman_4hrs_100ng | Test | 4hrs |
| SRR12291334 | Donor A | GSM4683213 | A11_caur_KTCbg_4hrs_100ng | Test | 4hrs |
| SRR12291335 | Donor A | GSM4683214 | A13_caur_KTClive_24hrs_100ng | Test | 24hrs |
| SRR12291336 | Donor A | GSM4683215 | A14_caur_KTCman_24hrs_100ng | Test | 24hrs |
| SRR12291337 | Donor A | GSM4683216 | A15_caur_KTCbg_24hrs_100ng | Test | 24hrs |
| SRR12291338 | Donor B | GSM4683217 | B1_RPMI_4hrs_250ng | Control | 4hrs |
| SRR12291342 | Donor B | GSM4683221 | B5_RPMI_24hrs_250ng | Control | 24hrs |
| SRR12291346 | Donor B | GSM4683225 | B9_caur_KTClive_4hrs_250ng | Test | 4hrs |
| SRR12291347 | Donor B | GSM4683226 | B10_caur_KTCman_4hrs_250ng | Test | 4hrs |
| SRR12291348 | Donor B | GSM4683227 | B11_caur_KTCbg_4hrs_250ng | Test | 4hrs |
| SRR12291349 | Donor B | GSM4683228 | B13_caur_KTClive_24hrs_250ng | Test | 24hrs |
| SRR12291350 | Donor B | GSM4683229 | B14_caur_KTCman_24hrs_250ng | Test | 24hrs |
| SRR12291351 | Donor B | GSM4683230 | B15_caur_KTCbg_24hrs_250ng | Test | 24hrs |
| SRR12291352 | Donor C | GSM4683231 | C1_RPMI_4hrs_250ng | Control | 4hrs |
| SRR12291356 | Donor C | GSM4683235 | C5_RPMI_24hrs_250ng | Control | 24hrs |
| SRR12291360 | Donor C | GSM4683239 | C9_caur_KTClive_4hrs_repl_250ng | Test | 4hrs |
| SRR12291361 | Donor C | GSM4683240 | C10_caur_KTCman_4hrs_250ng | Test | 4hrs |
| SRR12291362 | Donor C | GSM4683241 | C11_caur_KTCbg_4hrs_250ng | Test | 4hrs |
| SRR12291363 | Donor C | GSM4683242 | C12_caur_KTClive_24hrs_repl_250ng | Test | 24hrs |
| SRR12291364 | Donor C | GSM4683243 | C13_caur_KTCman_24hrs_250ng | Test | 24hrs |
| SRR12291365 | Donor C | GSM4683244 | C14_caur_KTCbg_24hrs_250ng | Test | 24hrs |

**supplementary table S1:** Raw RNA-sequencing data generated by *Bruno et al.* to investigate human immune responses to *Candida auris*. For this study we used 24 RNA-Seq reads data from NCBI BioProject PRJNA647871 representing three healthy human donors, four stimulation conditions, and two time points
