## Supplementary material for "Computational identification of candidate human long noncoding RNAs associated with PBMC responses to *Candida auris* and its cell wall components": Supplementary_Table_S3.docx (from another version of this preprint: version 1, 2026-09-22)

**Supplementary Table S3|** Class codes **u** and **x** constituted the primary novel-lncRNA discovery set. Class-code definitions follow GffCompare nomenclature.

| GffCompare class code | Transcript models |
| --- | --- |
| = | 384,464 |
| i | 6,703 |
| j | 3,772 |
| k | 2,457 |
| **x** | **1,294** |
| o | 481 |
| **u** | **388** |
| p | 273 |
| m | 138 |
| n | 121 |
| c | 103 |
| y | 10 |
| e | 1 |
| Total | 400,205 |
